## Supplementary Material for "Benchmarking 50 classification algorithms on 50 gene-expression datasets"

### Supplementary Figures

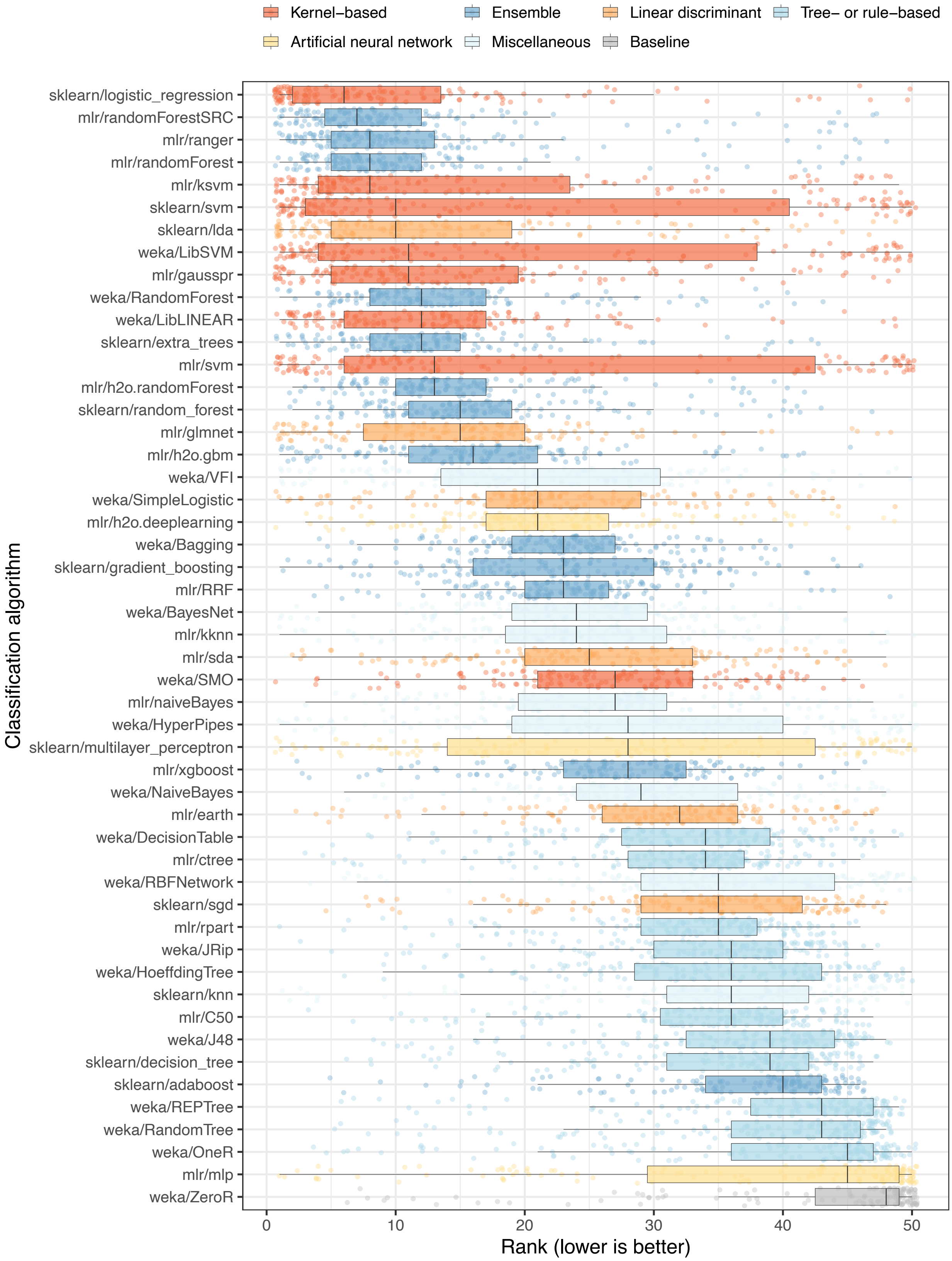

Kernel–based
Ensemble
Linear discriminant
Tree– or rule–based
Artificial neural network
Miscellaneous
Baseline
Classification algorithm
sklearn/logistic_regression
mlr/randomForestSRC
mlr/ranger
mlr/randomForest
mlr/ksvm
sklearn/svm
sklearn/lda
weka/LibSVM
mlr/gausspr
weka/RandomForest
weka/LibLINEAR
sklearn/extra_trees
mlr/svm
mlr/h2o.randomForest
sklearn/random_forest
mlr/glmnet
mlr/h2o.gbm
weka/VFI
weka/SimpleLogistic
mlr/h2o.deeplearning
weka/Bagging
sklearn/gradient_boosting
mlr/RRF
weka/BayesNet
mlr/kknn
mlr/sda
weka/SMO
mlr/naiveBayes
weka/HyperPipes
sklearn/multilayer_perceptron
mlr/xgboost
weka/NaiveBayes
mlr/earth
weka/DecisionTable
mlr/ctree
weka/RBFNetwork
sklearn/sgd
mlr/rpart
weka/JRip
weka/HoeffdingTree
sklearn/knn
mlr/C50
weka/J48
sklearn/decision_tree
sklearn/adaboost
weka/REPTree
weka/RandomTree
weka/OneR
mlr/mlp
weka/ZeroR
0
10
20
30
40
50
Rank (lower is better)


7

**Figure S1: Relative performance of classification algorithms using gene-expression predictors and area under the receiver operating characteristic curve as the metric.** We predicted patient states using gene-expression predictors only (Analysis 1). For each combination of dataset, class variable, and classification algorithm, we calculated the arithmetic mean of area under the receiver operating characteristic curve (AUROC) values across 50 iterations of Monte Carlo cross-validation. Next we sorted the algorithms based on the average rank across all dataset/class combinations. Each data point that overlays the box plots represents a particular dataset/class combination. The top 15 performers (relatively low ranks) were algorithms that use linear decision boundaries, kernel functions, and/or ensembles of decision trees.

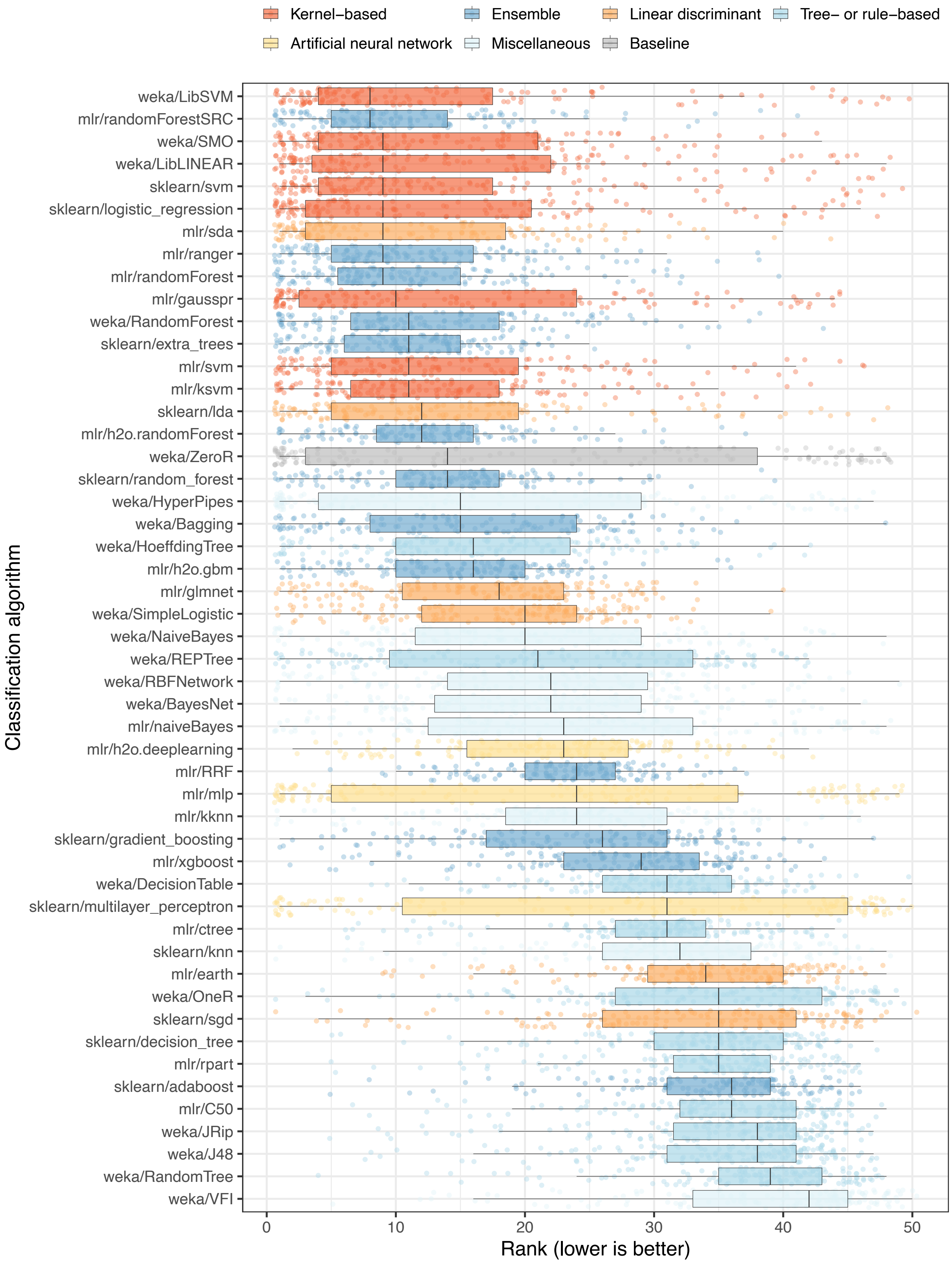

Kernel−based
Ensemble
Linear discriminant
Tree− or rule−based
Artificial neural network
Miscellaneous
Baseline
Classification algorithm
weka/LibSVM
mlr/randomForestSRC
weka/SMO
weka/LibLINEAR
sklearn/svm
sklearn/logistic_regression
mlr/sda
mlr/ranger
mlr/randomForest
mlr/gausspr
weka/RandomForest
sklearn/extra_trees
mlr/svm
mlr/ksvm
sklearn/lda
mlr/h2o.randomForest
weka/ZeroR
sklearn/random_forest
weka/HyperPipes
weka/Bagging
weka/HoeffdingTree
mlr/h2o.gbm
mlr/glmnet
weka/SimpleLogistic
weka/NaiveBayes
weka/REPTree
weka/RBFNetwork
weka/BayesNet
mlr/naiveBayes
mlr/h2o.deeplearning
mlr/RRF
mlr/mlp
mlr/kknn
sklearn/gradient_boosting
mlr/xgboost
weka/DecisionTable
sklearn/multilayer_perceptron
mlr/ctree
sklearn/knn
mlr/earth
weka/OneR
sklearn/sgd
sklearn/decision_tree
mlr/rpart
sklearn/adaboost
mlr/C50
weka/JRip
weka/J48
weka/RandomTree
weka/VFI
0
10
20
30
40
50
Rank (lower is better)


17

**Figure S2: Relative performance of classification algorithms using gene-expression predictors and classification accuracy as the metric.** We predicted patient states using gene-expression predictors only (Analysis 1). For each combination of dataset, class variable, and classification algorithm, we calculated the arithmetic mean of classification accuracy across 50 iterations of Monte Carlo cross-validation. Next we sorted the algorithms based on the average rank across all dataset/class combinations. Each data point that overlays the box plots represents a particular dataset/class combination.

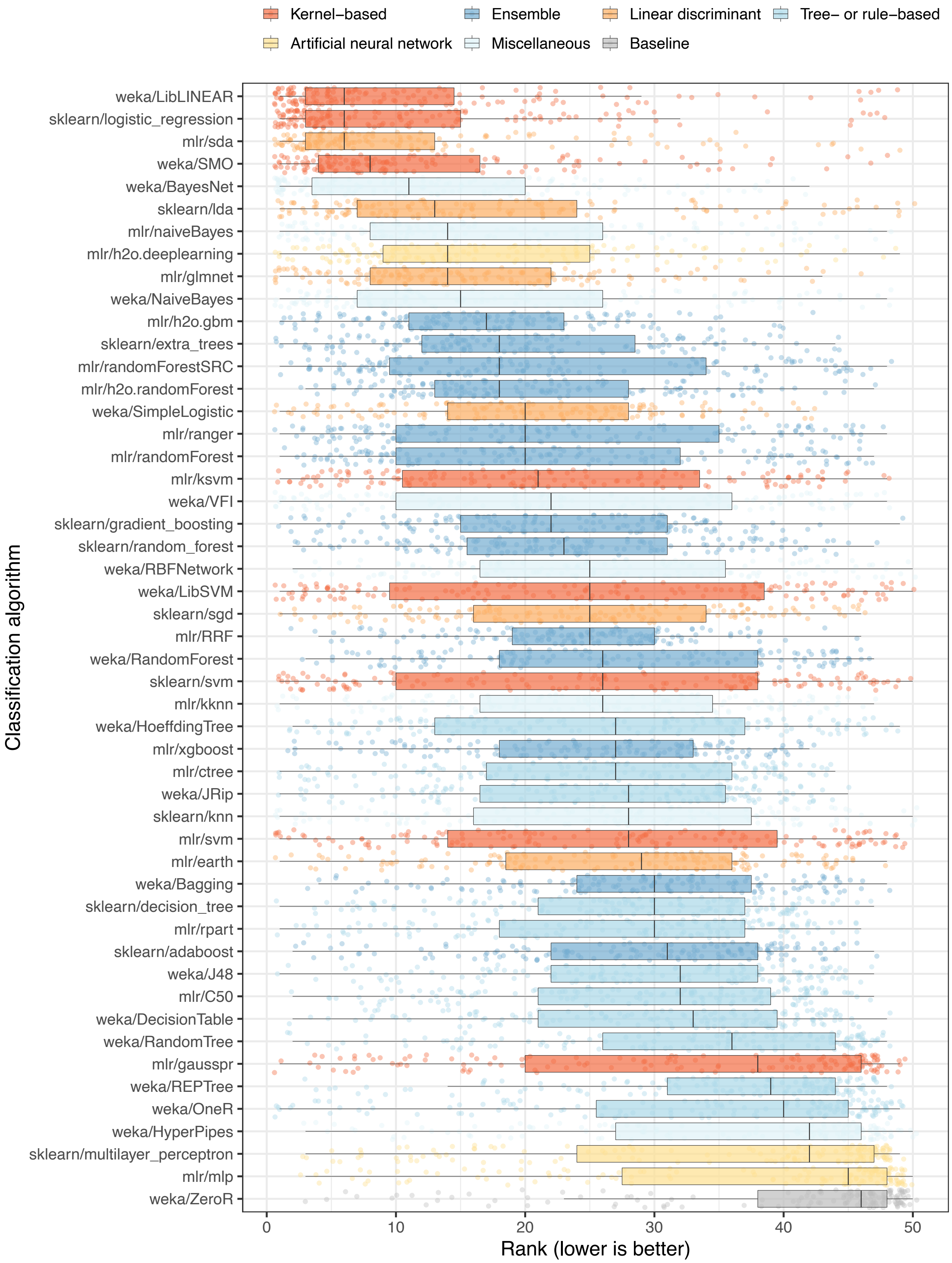

Kernel−based
Ensemble
Linear discriminant
Tree− or rule−based
Artificial neural network
Miscellaneous
Baseline
Classification algorithm
weka/LibLINEAR
sklearn/logistic_regression
mlr/sda
weka/SMO
weka/BayesNet
sklearn/lda
mlr/naiveBayes
mlr/h2o.deeplearning
mlr/glmnet
weka/NaiveBayes
mlr/h2o.gbm
sklearn/extra_trees
mlr/randomForestSRC
mlr/h2o.randomForest
weka/SimpleLogistic
mlr/ranger
mlr/randomForest
mlr/ksvm
weka/VFI
sklearn/gradient_boosting
sklearn/random_forest
weka/RBFNetwork
weka/LibSVM
sklearn/sgd
mlr/RRF
weka/RandomForest
sklearn/svm
mlr/kknn
weka/HoeffdingTree
mlr/xgboost
mlr/ctree
weka/JRip
sklearn/knn
mlr/svm
mlr/earth
weka/Bagging
sklearn/decision_tree
mlr/rpart
sklearn/adaboost
weka/J48
mlr/C50
weka/DecisionTable
weka/RandomTree
mlr/gausspr
weka/REPTree
weka/OneR
weka/HyperPipes
sklearn/multilayer_perceptron
mlr/mlp
weka/ZeroR
0
10
20
30
40
50
Rank (lower is better)


24

**Figure S3: Relative performance of classification algorithms using gene-expression predictors and Matthews Correlation Coefficient as the metric.** We predicted patient states using gene-expression predictors only (Analysis 1). For each combination of dataset, class variable, and classification algorithm, we calculated the arithmetic mean of the Matthews Correlation Coefficient across 50 iterations of Monte Carlo cross-validation. Next we sorted the algorithms based on the average rank across all dataset/class combinations. Each data point that overlays the box plots represents a particular dataset/class combination.

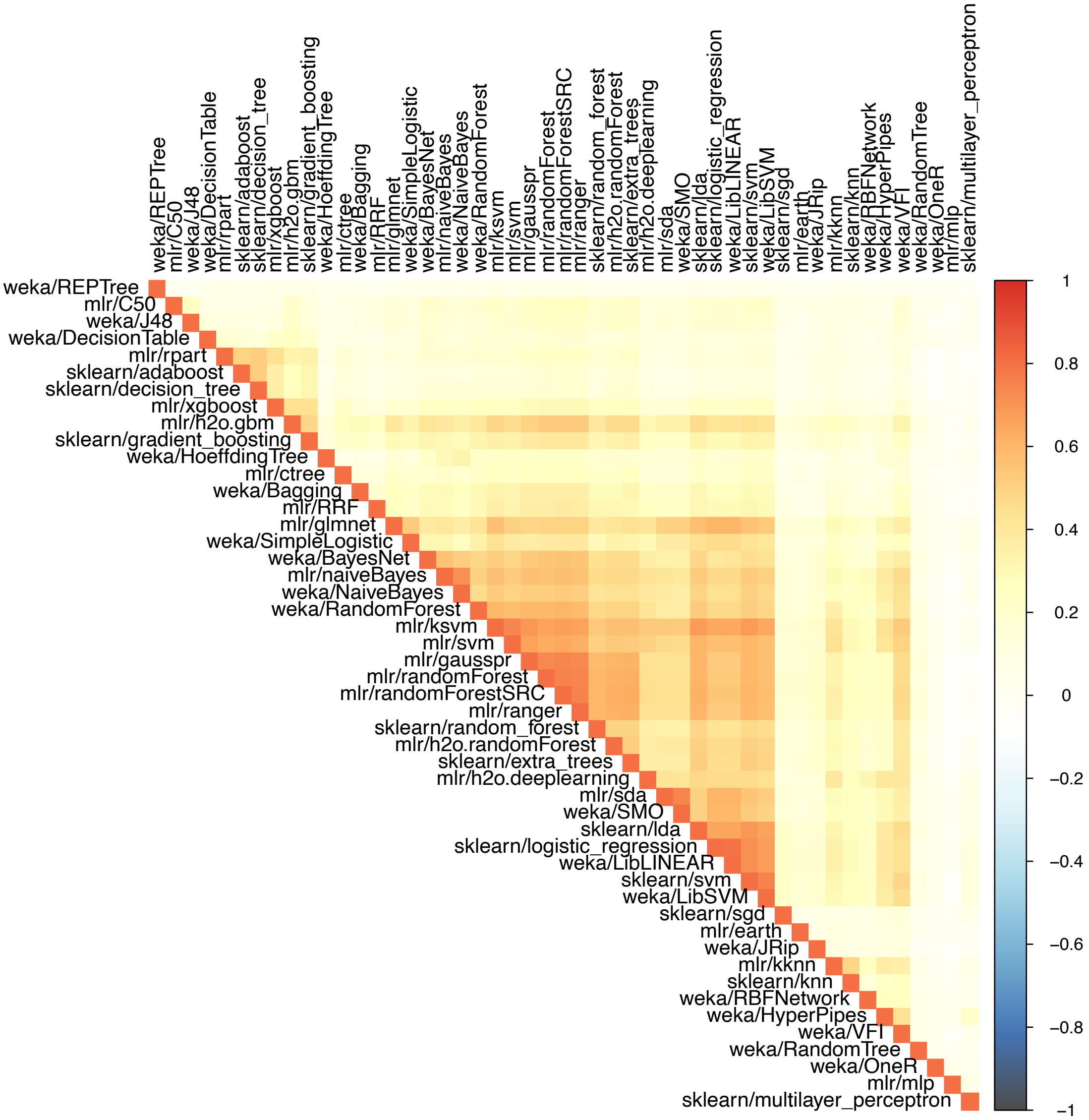


**Figure S4: Pairwise correlations of sample-level, probabilistic predictions between classification algorithms for dataset GSE10320.** We used each classification algorithm to make probabilistic predictions of relapse in Wilms tumor patients (GSE10320). Based on these predictions, we calculated the Spearman correlation coefficient for each pair of algorithms. These coefficients, averaged across Monte Carlo cross-validation iterations, are illustrated as a correlation plot, clustered based on similarity.

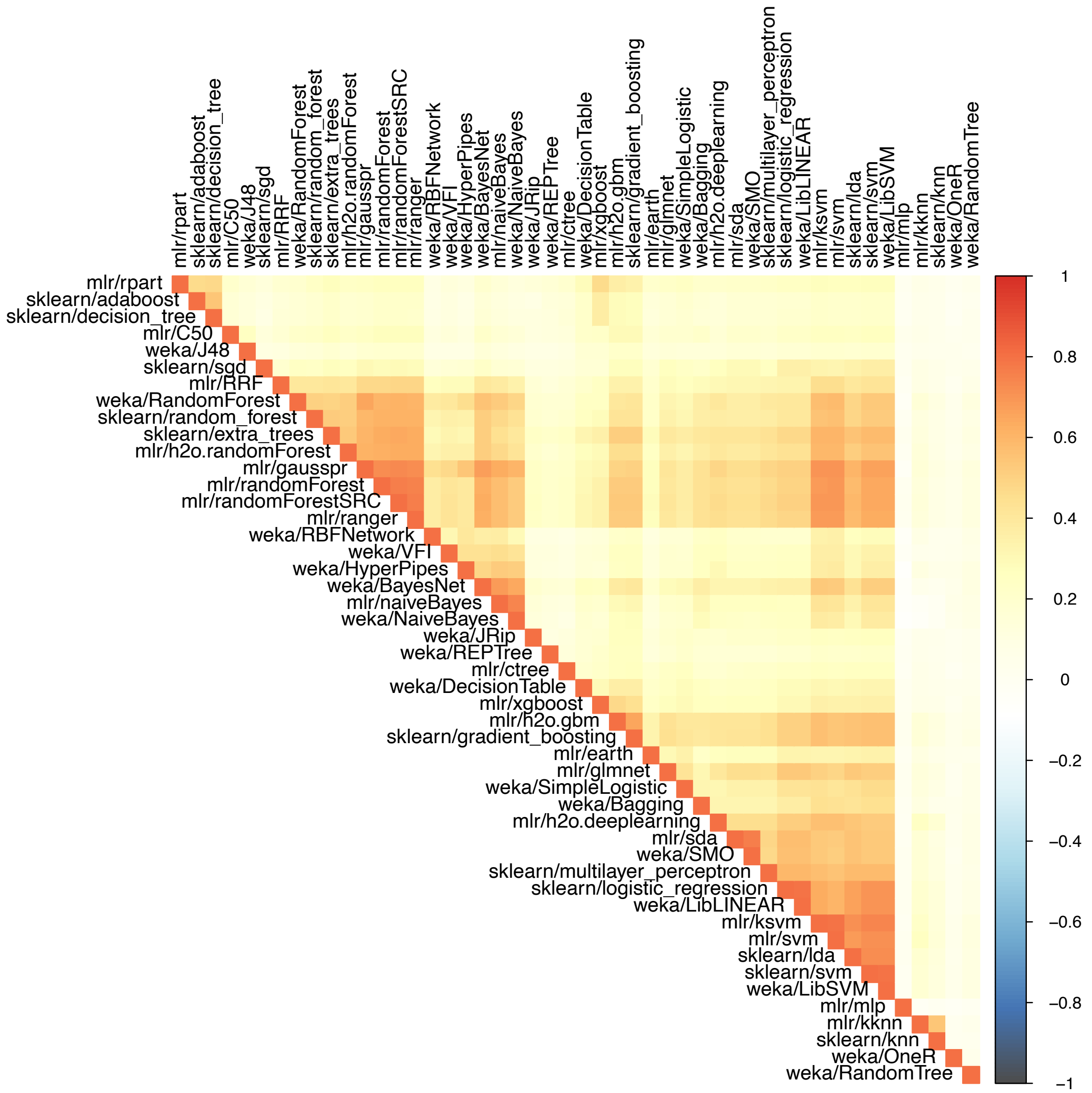


**Figure S5: Pairwise correlations of sample-level, probabilistic predictions between classification algorithms for dataset GSE46691.** We used each classification algorithm to make probabilistic predictions of early metastasis following radical prostatectomy (GSE46691). Based on these predictions, we calculated the Spearman correlation coefficient for each pair of algorithms. These coefficients, averaged across Monte Carlo cross-validation iterations, are illustrated as a correlation plot, clustered based on similarity.

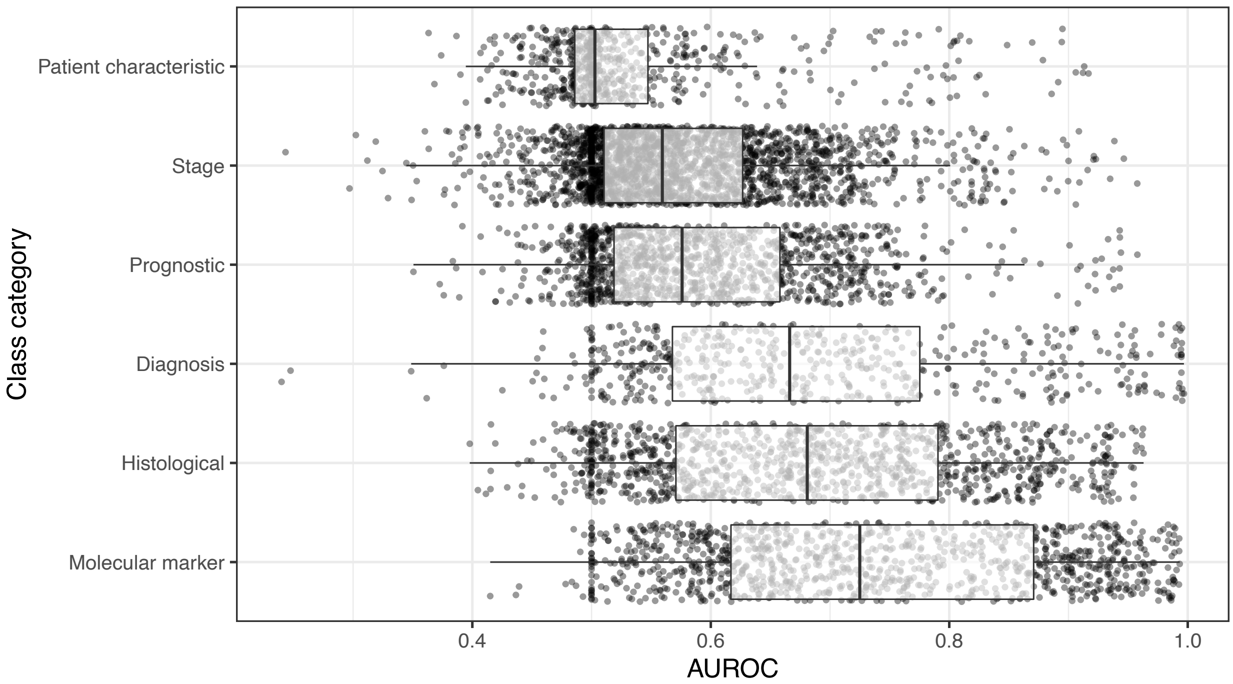


**Figure S6: Dataset performance by class category when using gene-expression predictors.** For each class variable across all datasets, we assigned a category representing the type of patient state being predicted. For Analysis 1, we show the predictive performance for each combination of dataset, class variable, and classification algorithm in each class category. We use area under the receiver operating characteristic curve (AUROC) as the metric. The dashed, red line indicates the performance expected by random chance. The top-performing category was “Molecular Marker,” which includes class variables associated with mutation status, immunohistochemistry markers of protein expression, presence or absence of chromosomal aberrations, etc. The lowest-performing category was “Patient Characteristic,” which includes variables that indicate whether patients had a family history of cancer, had been diagnosed with multiple tumors, patient performance status, etc.

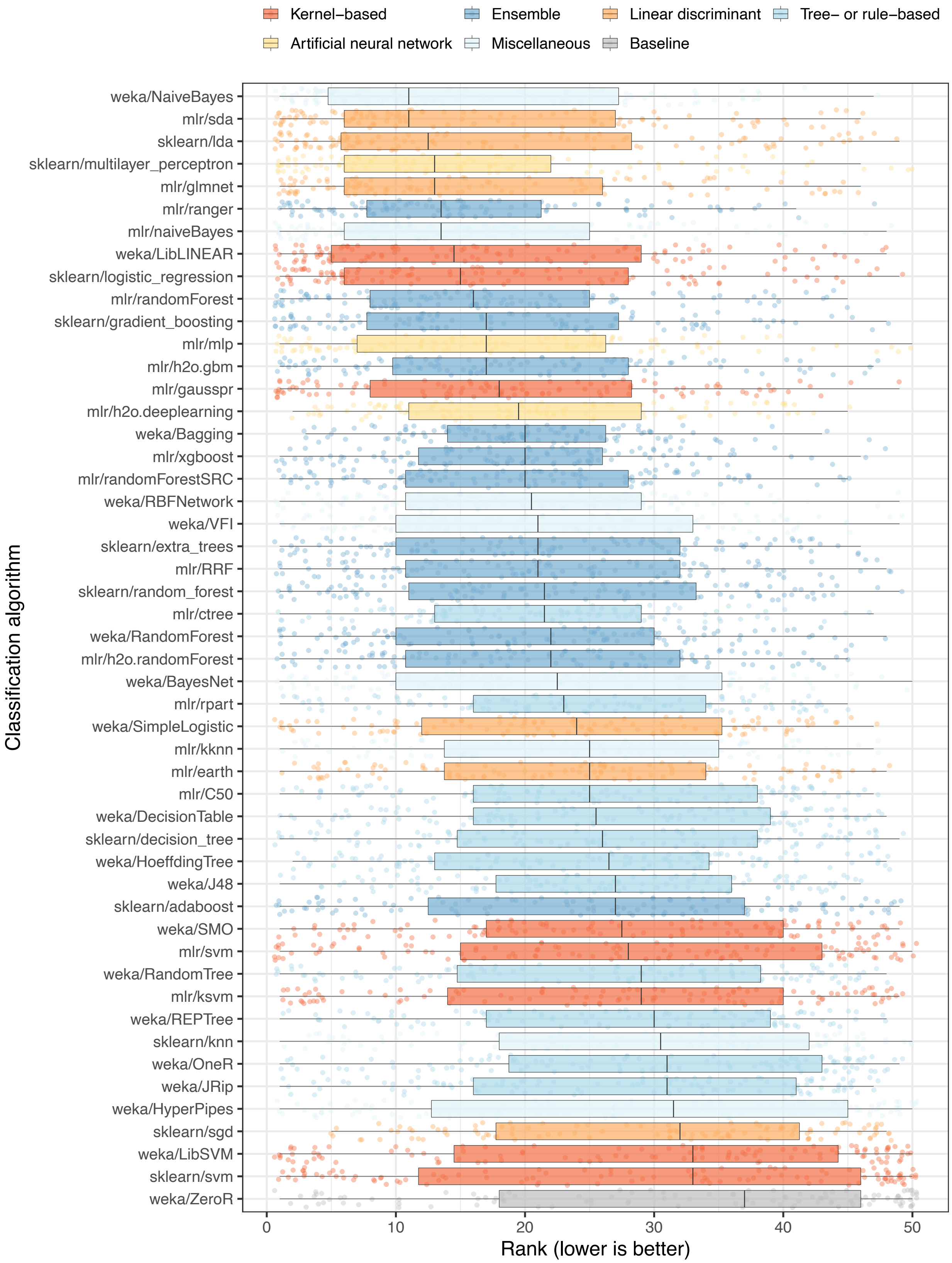

Kernel−based
Ensemble
Linear discriminant
Tree− or rule−based
Artificial neural network
Miscellaneous
Baseline
weka/NaiveBayes
mlr/sda
sklearn/lda
sklearn/multilayer_perceptron
mlr/glmnet
mlr/ranger
mlr/naiveBayes
weka/LibLINEAR
sklearn/logistic_regression
mlr/randomForest
sklearn/gradient_boosting
mlr/mlp
mlr/h2o.gbm
mlr/gausspr
mlr/h2o.deeplearning
weka/Bagging
mlr/xgboost
mlr/randomForestSRC
weka/RBFNetwork
weka/VFI
sklearn/extra_trees
mlr/RRF
sklearn/random_forest
mlr/ctree
weka/RandomForest
mlr/h2o.randomForest
weka/BayesNet
mlr/rpart
weka/SimpleLogistic
mlr/kknn
mlr/earth
mlr/C50
weka/DecisionTable
sklearn/decision_tree
weka/HoeffdingTree
weka/J48
sklearn/adaboost
weka/SMO
mlr/svm
weka/RandomTree
mlr/ksvm
weka/REPTree
sklearn/knn
weka/OneR
weka/JRip
weka/HyperPipes
sklearn/sgd
weka/LibSVM
sklearn/svm
weka/ZeroR
Classification algorithm
0
10
20
30
40
50
Rank (lower is better)


56

**Figure S7: Relative performance of classification algorithms using clinical predictors and area under the receiver operating characteristic curve as the metric.** We predicted patient states using clinical predictors only (Analysis 2). For each combination of dataset, class variable, and classification algorithm, we calculated the arithmetic mean of area under the receiver operating characteristic curve (AUROC) values across 50 iterations of Monte Carlo cross-validation. Next we sorted the algorithms based on the average rank across all dataset/class combinations. Each data point that overlays the box plots represents a particular dataset/class combination (some datasets did not have clinical predictors). The top-performing algorithms (relatively low ranks) were similar overall to Analysis 1; however, some differences were large. For example, `weka/NaiveBayes` performed best overall in Analysis 2 but was ranked 28th in Analysis 1.

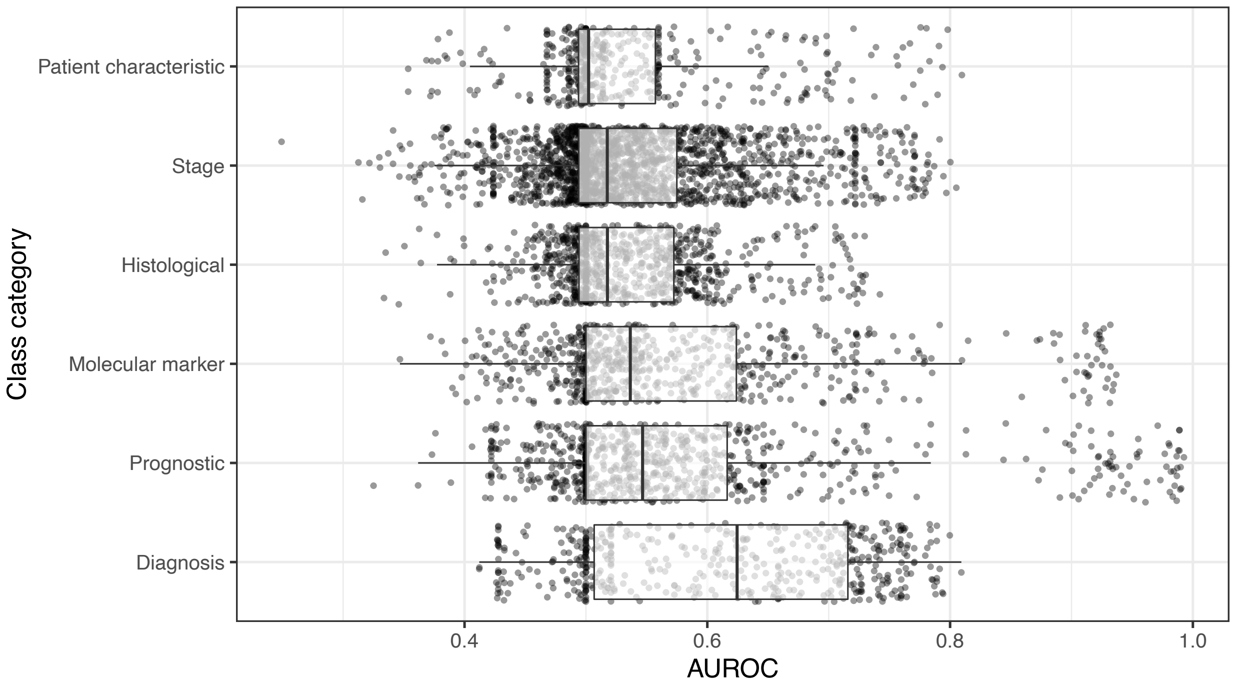


**Figure S8: Dataset performance by class category when using clinical predictors.** For each class variable across all datasets, we assigned a category representing the type of patient state being predicted. For Analysis 2, we show the predictive performance for each combination of dataset, class variable, and classification algorithm in each class category. We use area under the receiver operating characteristic curve (AUROC) as the metric. The dashed, red line indicates the performance expected by random chance. The top-performing category was "Diagnosis," which includes class variables associated with a particular disease or subtype. The lowest-performing category was "Patient Characteristic," which includes variables that indicate whether patients had a family history of cancer, had been diagnosed with multiple tumors, patient performance status, etc.

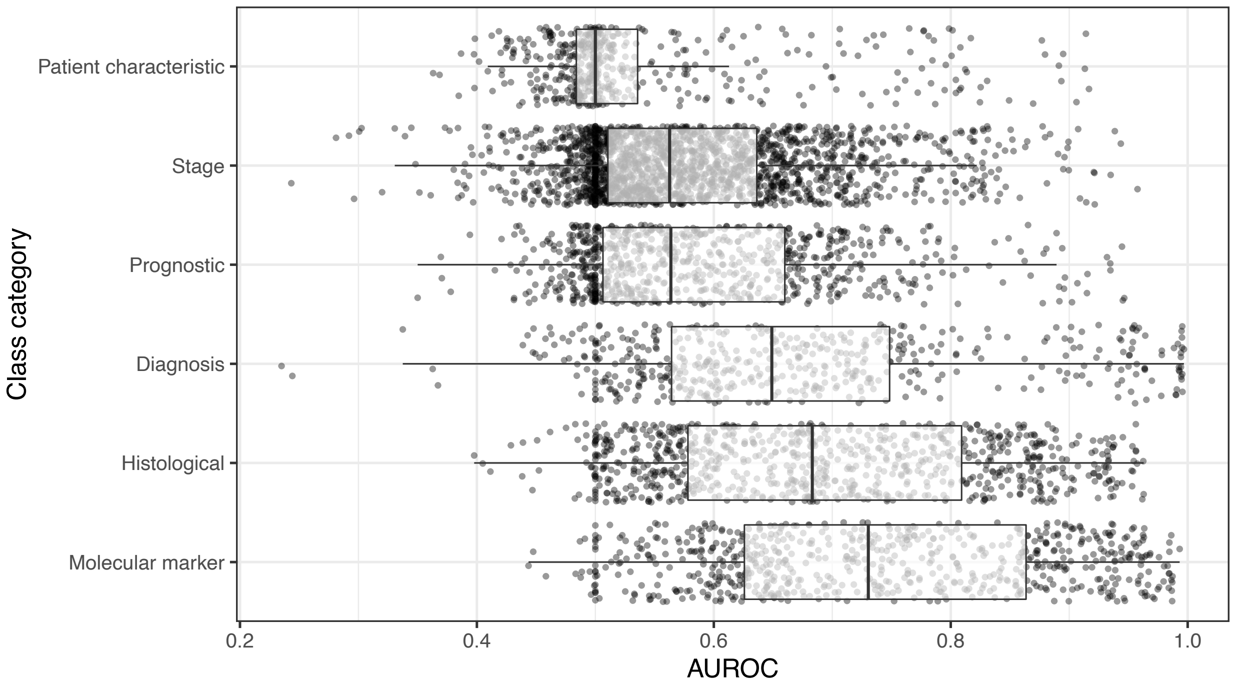


**Figure S9: Dataset performance by class category when using gene-expression and clinical predictors.** For each class variable across all datasets, we assigned a category representing the type of patient state being predicted. For Analysis 3, we show the predictive performance for each combination of dataset, class variable, and classification algorithm in each class category. We use area under the receiver operating characteristic curve (AUROC) as a metric. The dashed, red line indicates the performance expected by random chance. As with Analysis 1 (Figure S6), the top-performing category was "Molecular Marker," which includes class variables associated with mutation status, immunohistochemistry markers of protein expression, presence or absence of chromosomal aberrations, etc. The lowest-performing category was "Patient Characteristic," which includes variables that indicate whether patients had a family history of cancer, had been diagnosed with multiple tumors, patient performance status, etc.

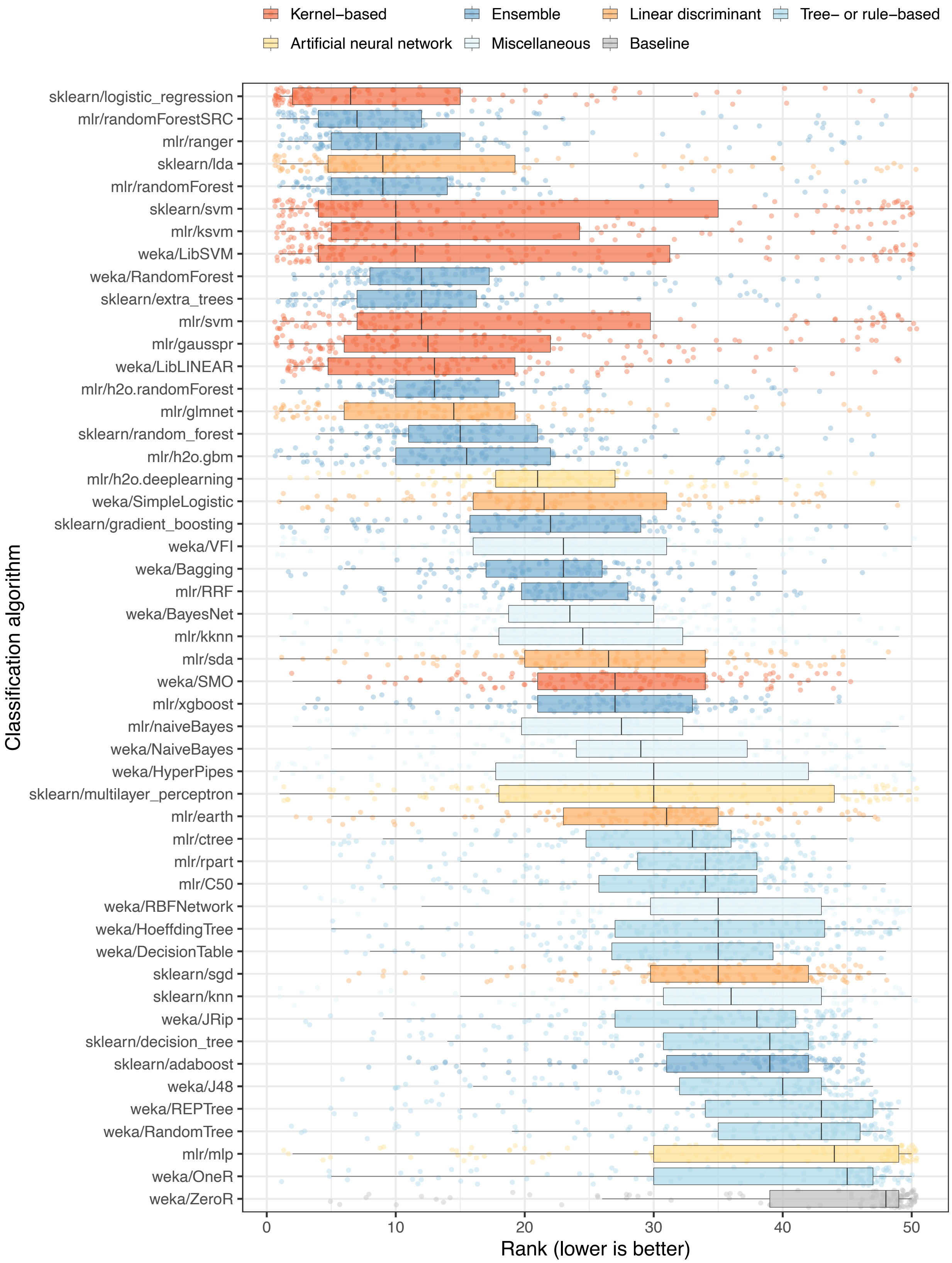
Kernel–based
Ensemble
Linear discriminant
Tree– or rule–based
Artificial neural network
Miscellaneous
Baseline
Classification algorithm
sklearn/logistic_regression
mlr/randomForestSRC
mlr/ranger
sklearn/lda
mlr/randomForest
sklearn/svm
mlr/ksvm
weka/LibSVM
weka/RandomForest
sklearn/extra_trees
mlr/svm
mlr/gausspr
weka/LibLINEAR
mlr/h2o.randomForest
mlr/glmnet
sklearn/random_forest
mlr/h2o.gbm
mlr/h2o.deeplearning
weka/SimpleLogistic
sklearn/gradient_boosting
weka/VFI
weka/Bagging
mlr/RRF
weka/BayesNet
mlr/kknn
mlr/sda
weka/SMO
mlr/xgboost
mlr/naiveBayes
weka/NaiveBayes
weka/HyperPipes
sklearn/multilayer_perceptron
mlr/earth
mlr/ctree
mlr/rpart
mlr/C50
weka/RBFNetwork
weka/HoeffdingTree
weka/DecisionTable
sklearn/sgd
sklearn/knn
weka/JRip
sklearn/decision_tree
sklearn/adaboost
weka/J48
weka/REPTree
weka/RandomTree
mlr/mlp
weka/OneR
weka/ZeroR
0
10
20
30
40
50
Rank (lower is better)

89

**Figure S10: Relative performance of classification algorithms using gene-expression and clinical predictors** We predicted patient states using gene-expression and clinical predictors (Analysis 3). For each combination of dataset, class variable, and classification algorithm, we calculated the arithmetic mean of area under the receiver operating characteristic curve (AUROC) values across 50 iterations of Monte Carlo cross-validation. Next we sorted the algorithms based on the average rank across all dataset/class combinations. Each data point that overlays the box plots represents a particular dataset/class combination. The algorithm ranks were highly similar to the ranks for Analysis 1 (Figure S1).

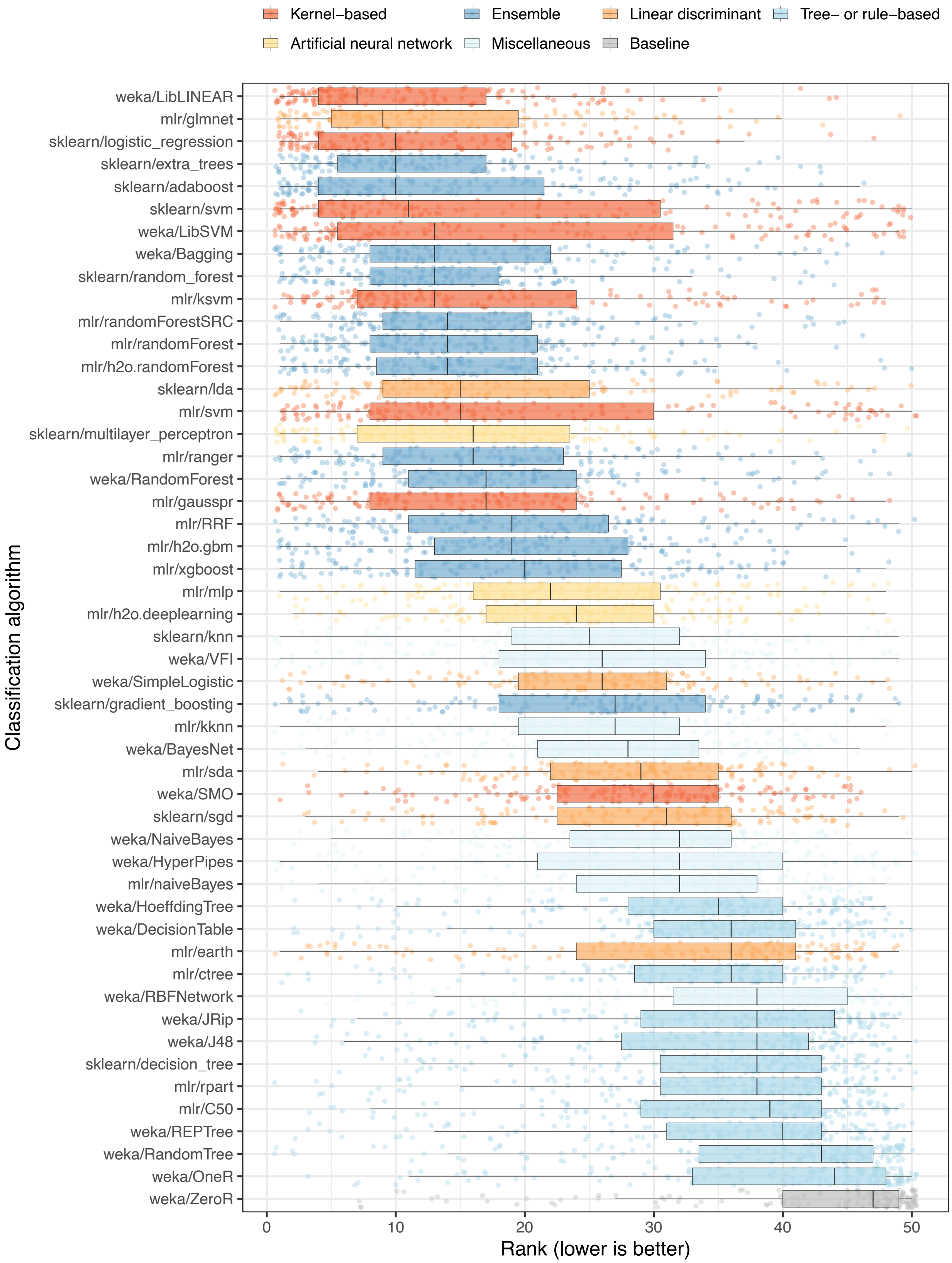
Kernel−based
Ensemble
Linear discriminant
Tree− or rule−based
Artificial neural network
Miscellaneous
Baseline
Classification algorithm
weka/LibLINEAR
mlr/glmnet
sklearn/logistic_regression
sklearn/extra_trees
sklearn/adaboost
sklearn/svm
weka/LibSVM
weka/Bagging
sklearn/random_forest
mlr/ksvm
mlr/randomForestSRC
mlr/randomForest
mlr/h2o.randomForest
sklearn/lda
mlr/svm
sklearn/multilayer_perceptron
mlr/ranger
weka/RandomForest
mlr/gausspr
mlr/RRF
mlr/h2o.gbm
mlr/xgboost
mlr/mlp
mlr/h2o.deeplearning
sklearn/knn
weka/VFI
weka/SimpleLogistic
sklearn/gradient_boosting
mlr/kknn
weka/BayesNet
mlr/sda
weka/SMO
sklearn/sgd
weka/NaiveBayes
weka/HyperPipes
mlr/naiveBayes
weka/HoeffdingTree
weka/DecisionTable
mlr/earth
mlr/ctree
weka/RBFNetwork
weka/JRip
weka/J48
sklearn/decision_tree
mlr/rpart
mlr/C50
weka/REPTree
weka/RandomTree
weka/OneR
weka/ZeroR
0
10
20
30
40
50
Rank (lower is better)

97

**Figure S11: Relative performance of classification algorithms using gene-expression and clinical predictors and performing hyperparameter optimization.** We predicted patient states using gene-expression and clinical predictors with hyperparameter optimization (Analysis 4). We used nested cross validation to estimate which hyperparameter combination would be optimal for each algorithm in each training set. For each combination of dataset, class variable, and classification algorithm, we calculated the arithmetic mean of area under the receiver operating characteristic curve (AUROC) values across 5 iterations of Monte Carlo cross-validation. Next we sorted the algorithms based on the average rank across all dataset/class combinations. Each data point that overlays the box plots represents a particular dataset/class combination. The algorithm rankings followed similar trends as Analysis 3 (no hyperparameter optimization); however, some differences are notable. For example, the `weka/LibLINEAR` and `mlr/glmnet` algorithms were ranked 11th and 16th in Analysis 3 (Figure S10), but they were ranked 1st and 2nd in this analysis.

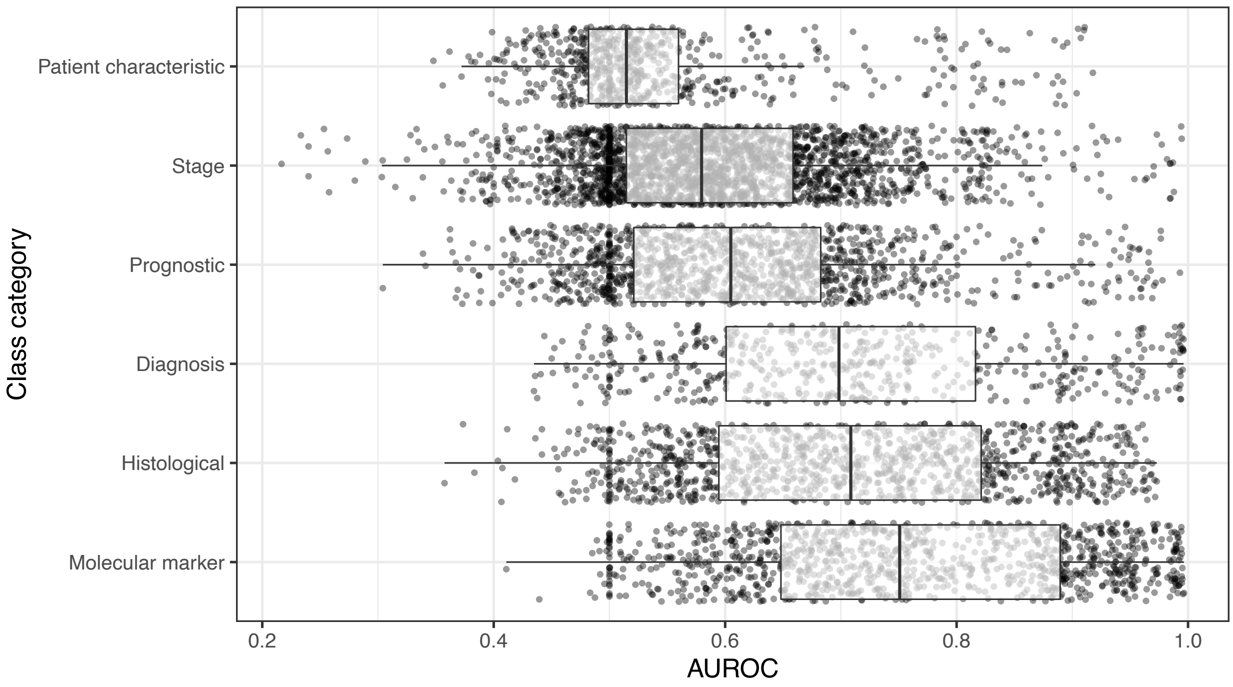


**Figure S12: Dataset performance by class category when using gene-expression and clinical predictors and performing hyperparameter optimization.** For each class variable across all datasets, we assigned a category representing the type of patient state being predicted. For Analysis 4, we show the predictive performance for each combination of dataset, class variable, and classification algorithm in each class category. We use area under the receiver operating characteristic curve (AUROC) as a metric. The dashed, red line indicates the performance expected by random chance. The results are similar to those of Analysis 3 (Figure S9).

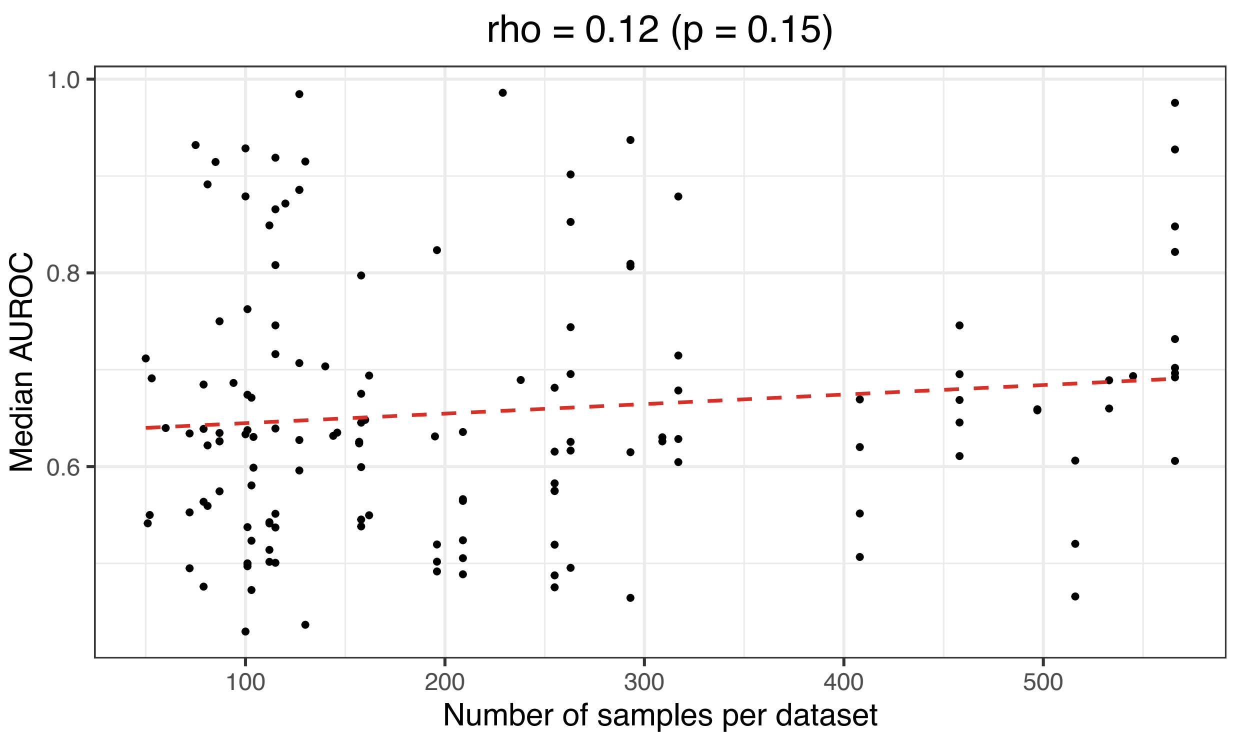


**Figure S13: Relationship between predictive performance and number of samples per dataset.** The number of patient samples differed by dataset. This scatterplot shows the relationship between the median area under the receiver operating characteristic curve (AUROC) and the number of samples in each dataset. We did not observe a significant relationship between these variables.

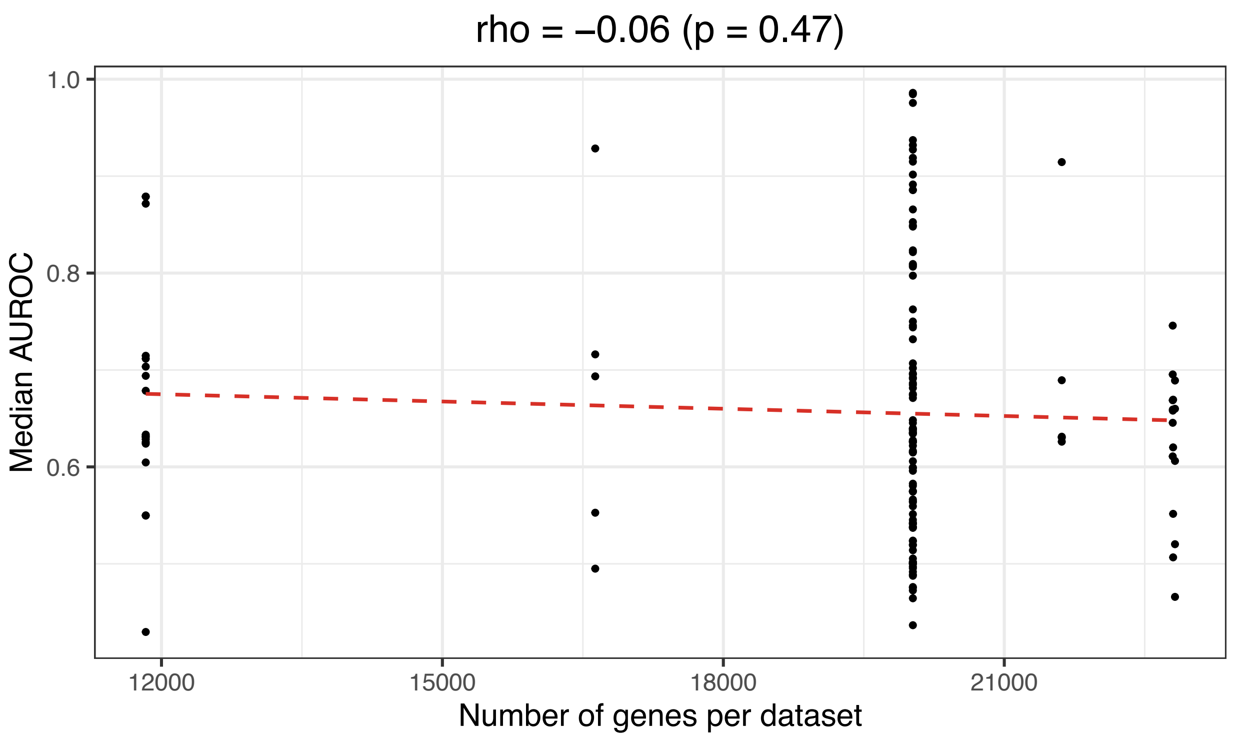


**Figure S14: Relationship between predictive performance and number of genes per dataset.** Due to differences in gene-expression profiling platforms, we had data for more genes in some datasets than in others. This scatterplot shows the relationship between the median area under the receiver operating characteristic curve (AUROC) and the number of genes in each dataset. We did not observe a significant relationship between these variables.

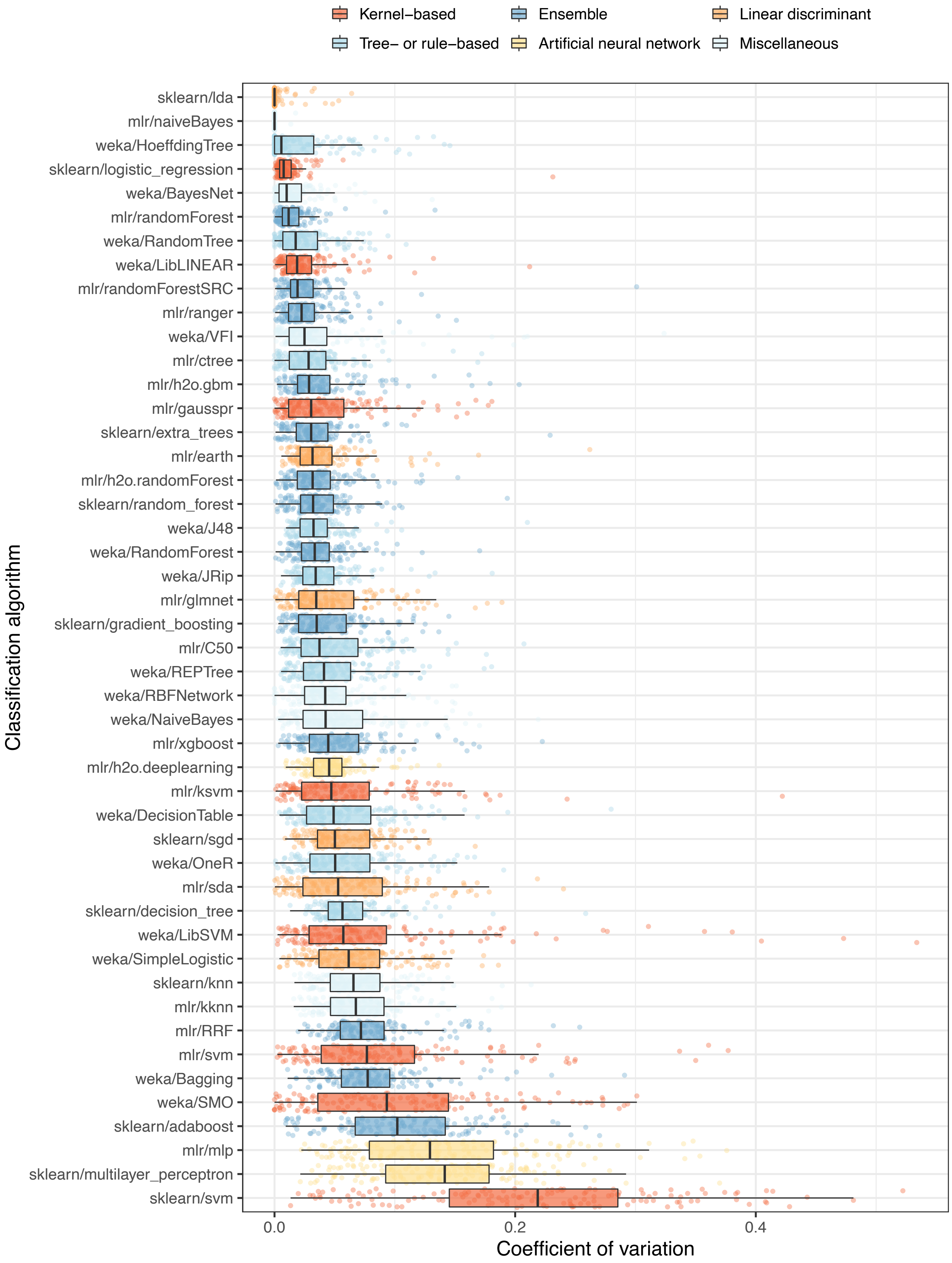
Kernel−based
Ensemble
Linear discriminant
Tree− or rule−based
Artificial neural network
Miscellaneous
Classification algorithm
sklearn/lda
mlr/naiveBayes
weka/HoeffdingTree
sklearn/logistic_regression
weka/BayesNet
mlr/randomForest
weka/RandomTree
weka/LibLINEAR
mlr/randomForestSRC
mlr/ranger
weka/VFI
mlr/ctree
mlr/h2o.gbm
mlr/gausspr
sklearn/extra_trees
mlr/earth
mlr/h2o.randomForest
sklearn/random_forest
weka/J48
weka/RandomForest
weka/JRip
mlr/glmnet
sklearn/gradient_boosting
mlr/C50
weka/REPTree
weka/RBFNetwork
weka/NaiveBayes
mlr/xgboost
mlr/h2o.deeplearning
mlr/ksvm
weka/DecisionTable
sklearn/sgd
weka/OneR
mlr/sda
sklearn/decision_tree
weka/LibSVM
weka/SimpleLogistic
sklearn/knn
mlr/kknn
mlr/RRF
mlr/svm
weka/Bagging
weka/SMO
sklearn/adaboost
mlr/mlp
sklearn/multilayer_perceptron
sklearn/svm
0.0
0.2
0.4
Coefficient of variation

130

**Figure S15: Variation in predictive performance across hyperparameter combinations.** In Analysis 4, we used nested cross validation to evaluate multiple hyperparameter combinations for each classification algorithm. We assessed the extent to which the area under the receiver operating characteristic curve (AUROC) varied across the hyperparameter combinations for each algorithm. For each combination of dataset, class variable, classification algorithm, and hyperparameter set, we averaged AUROC values across 5 Monte Carlo cross-validation iterations. Then we calculated the coefficient of variation for these averaged values across each combination of dataset/class and classification algorithm. Relatively low values indicate that the hyperparameter sets resulted in similar predictive performance. No results are available for 3 algorithms that used only a single hyperparameter option.

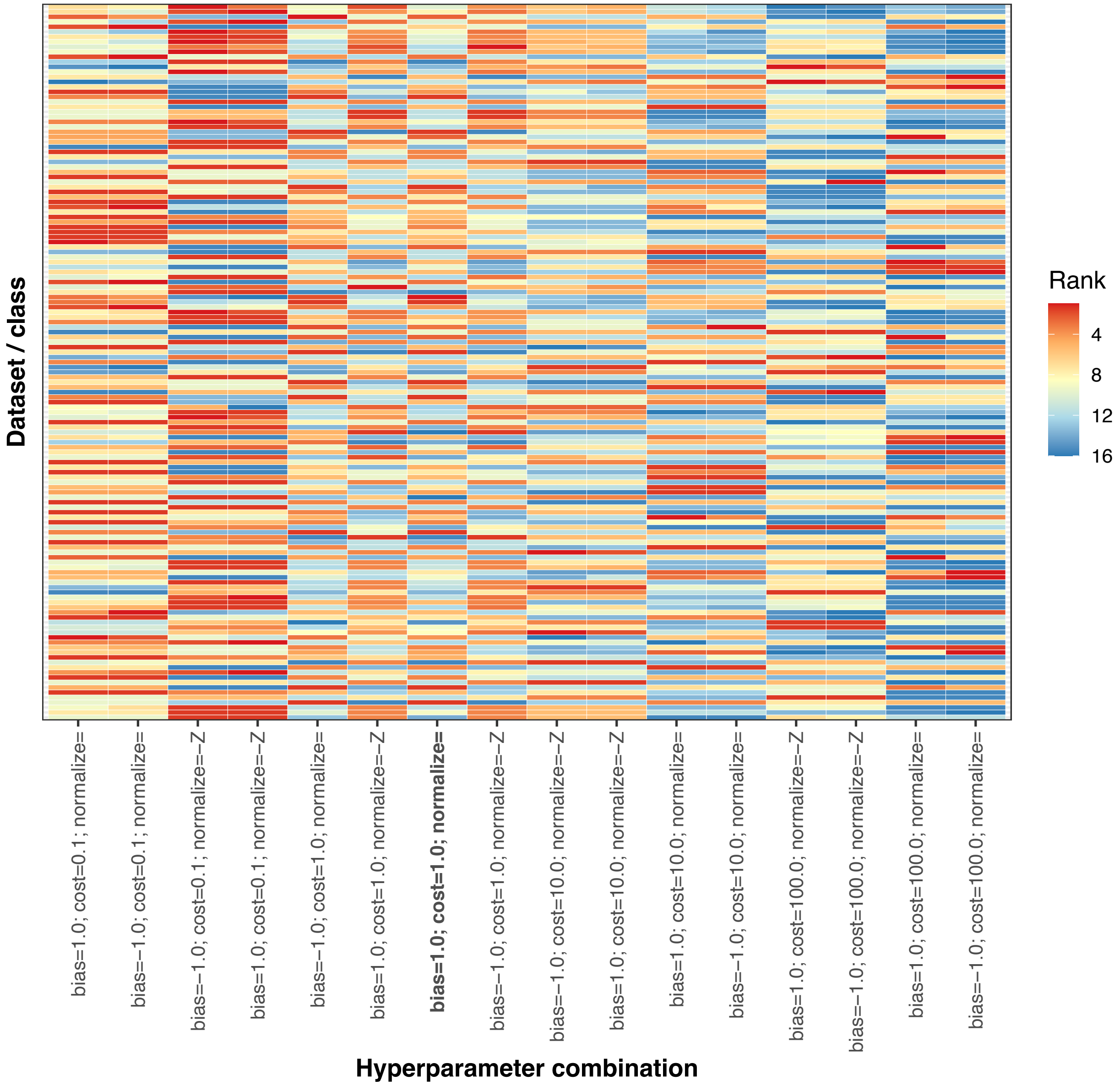


**Figure S16: Relative performance of different hyperparameter combinations for the `weka/LIBLINEAR` classification algorithm.** The ShinyLearner software supports 16 hyperparameter combinations for the `weka/LIBLINEAR` classification algorithm. In Analysis 4, we used nested cross validation for hyperparameter optimization. For each combination of dataset and class variable, we averaged the area under the receiver operating characteristic curve (AUROC) across all (outer) Monte Carlo cross-validation iterations and then ranked the averages for each hyperparameter combination.

Some combinations consistently outperformed other combinations, and the default combination performed suboptimally. Using relatively small `cost` values appeared to improve the performance more than any other option. This hyperparameter controls the regularization strength.

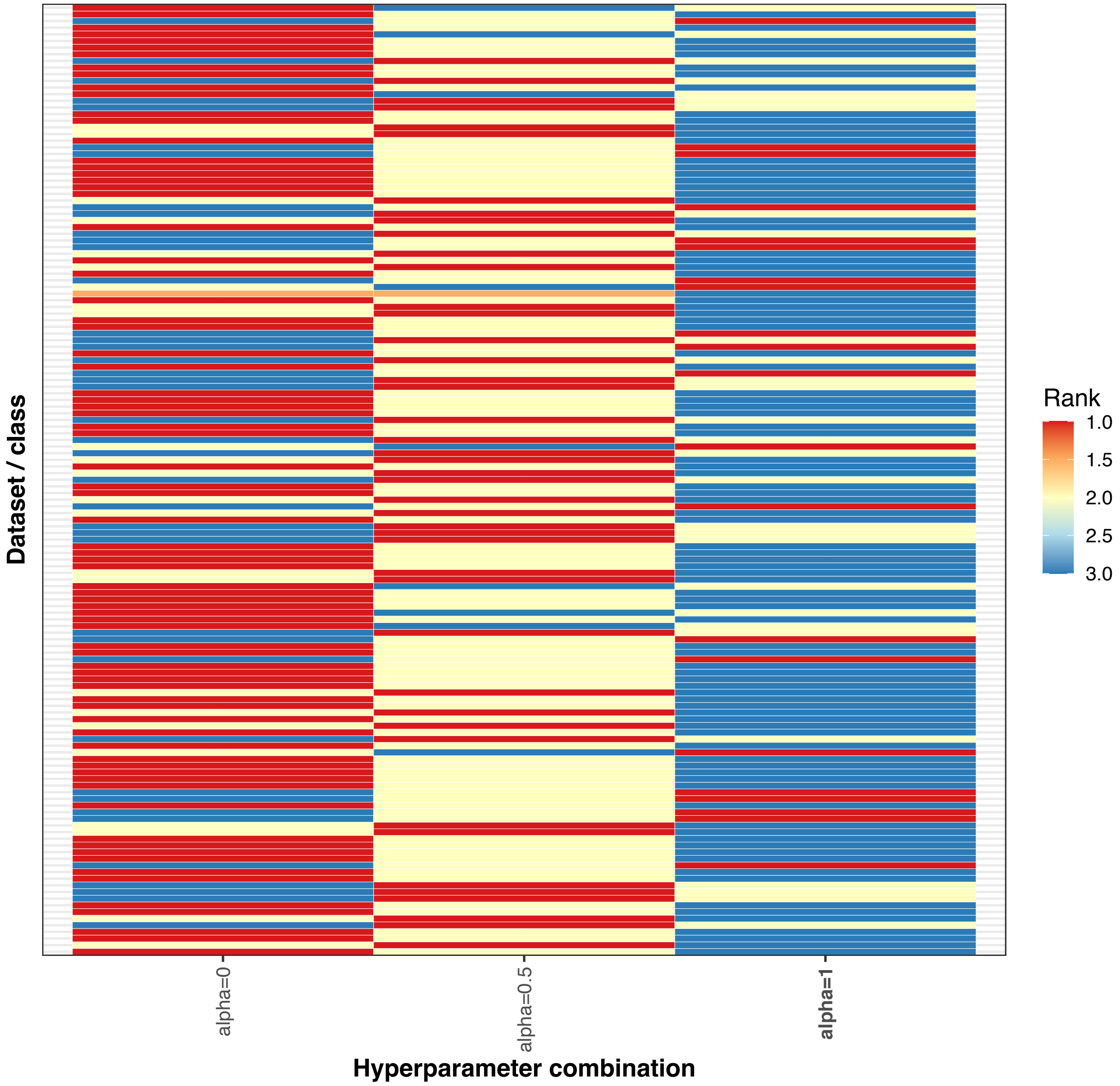


**Figure S17: Relative performance of different hyperparameter combinations for the `mlr/glmnet` classification algorithm.** The ShinyLearner software supports 3 hyperparameter combinations for the `mlr/glmnet` classification algorithm. In Analysis 4, we used nested cross validation for hyperparameter optimization. For each combination of dataset and class variable, we averaged the area under the receiver operating characteristic curve (AUROC) across all (outer) Monte Carlo cross-validation iterations and then ranked the averages for each hyperparameter combination. Using an `alpha` value of 0.5 or 0 resulted in better performance than a value of 1.

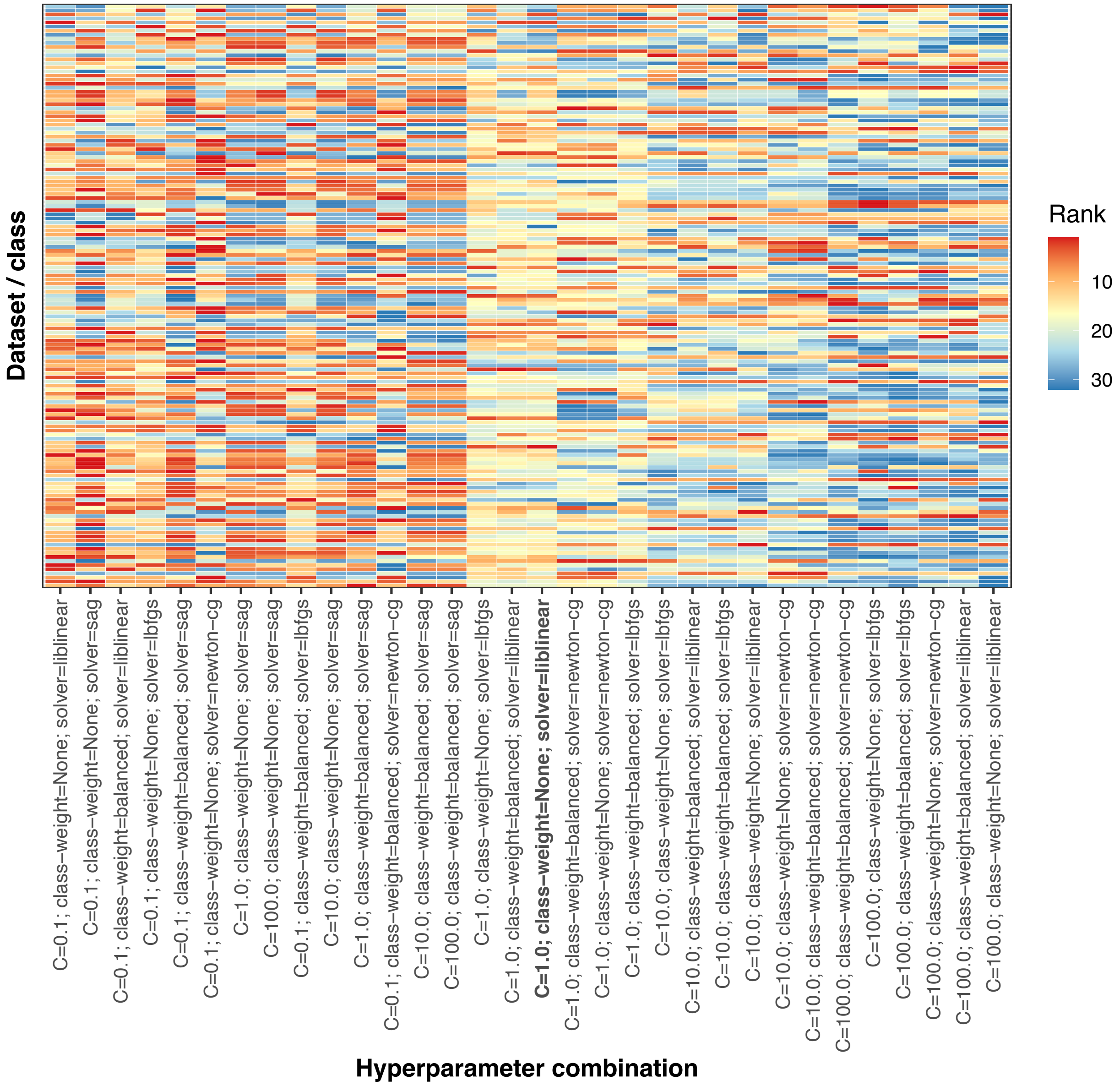


**Figure S18: Relative performance of different hyperparameter combinations for the `sklearn/logistic_regression` classification algorithm.** The ShinyLearner software supports 32 hyperparameter combinations for the `sklearn/logistic_regression` classification algorithm. In Analysis 4, we used nested cross validation for hyperparameter optimization. For each combination of dataset and class variable, we averaged the area under the receiver operating characteristic curve (AUROC) across all (outer) Monte Carlo cross-validation iterations and then ranked the averages for each hyperparameter combination. Some combinations consistently outperformed other combinations, and the default combination performed suboptimally. Using relatively small `cost` values appeared to improve the performance more than any other option. This hyperparameter controls the regularization strength.

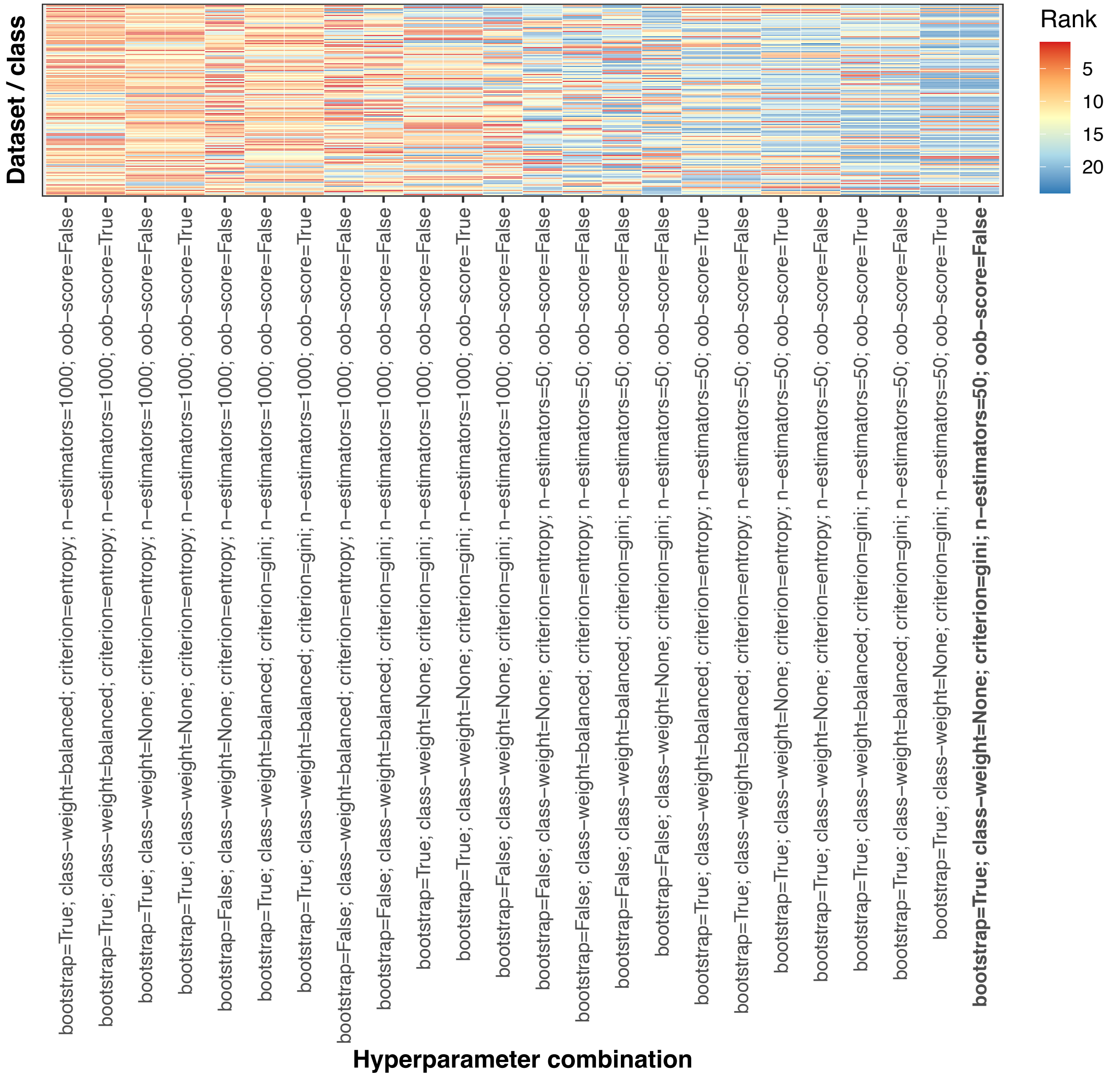


**Figure S19: Relative performance of different hyperparameter combinations for the `sklearn/extra_trees` classification algorithm.** The ShinyLearner software supports 24 hyperparameter combinations for the `sklearn/extra_trees` classification algorithm. In Analysis 4, we used nested cross validation for hyperparameter optimization. For each combination of dataset and class variable, we averaged the area under the receiver operating characteristic curve (AUROC) across all (outer) Monte Carlo cross-validation iterations and then ranked the averages for each hyperparameter combination. Some combinations consistently outperformed other combinations, and the default combination performed suboptimally. Using a larger number (n = 1000) of estimators (trees) appeared to improve the performance more than any other option.

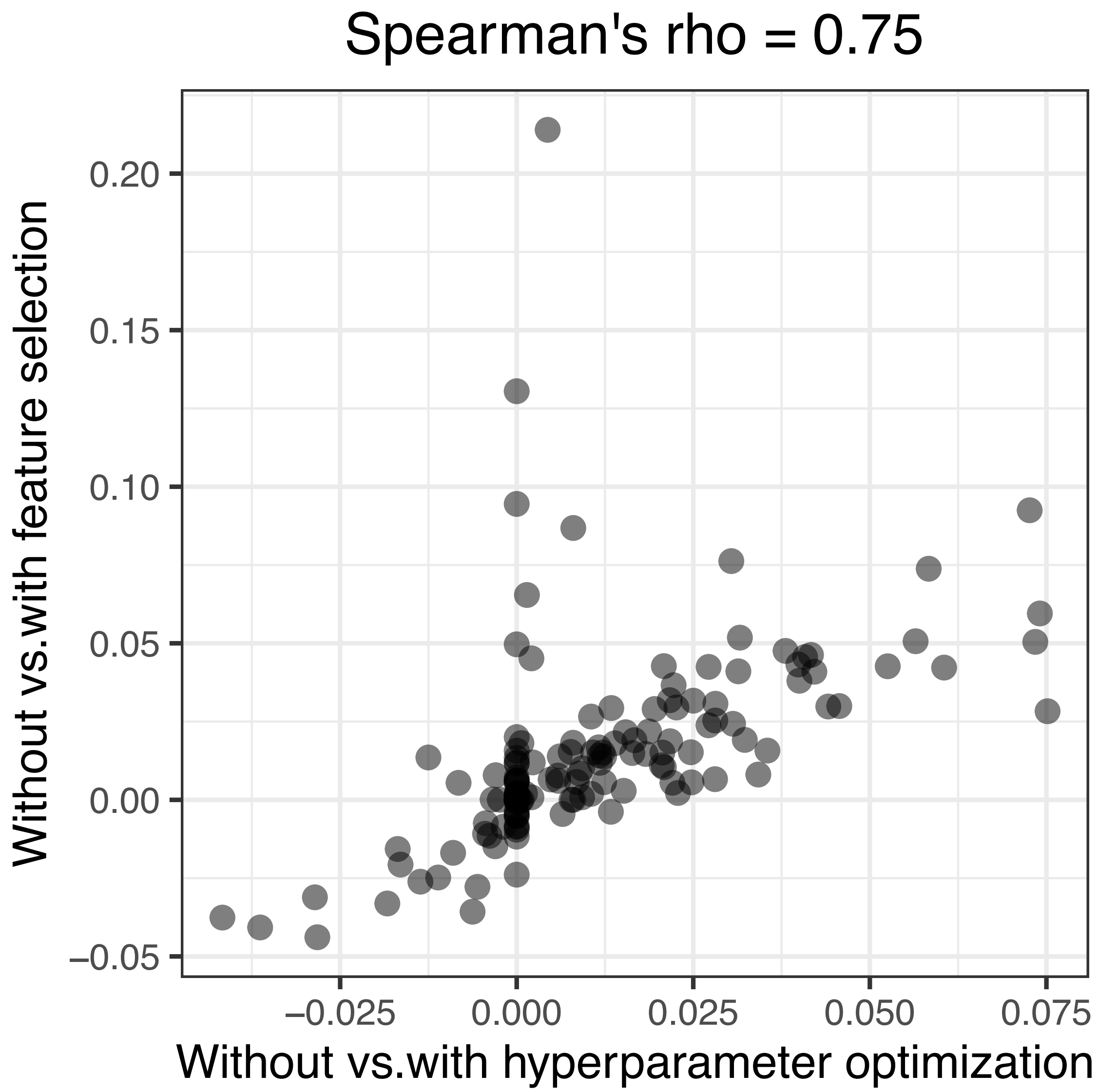


**Figure S20: Relative predictive performance when using hyperparameter optimization vs. feature selection.** We use as a baseline the predictive performance that we attained using default hyperparameters for the classification algorithms (Analysis 3). We quantified predictive performance using the area under the receiver operating characteristic curve (AUROC). This graph shows the increase or decrease in performance when selecting hyperparameters or selecting features relative to the baseline. Each point represents a particular combination of dataset and class variable. Generally, the dataset/class combinations that benefitted from hyperparameter optimization also benefitted from feature selection.

However, some dataset/class combinations that did not benefit from hyperparameter optimization did benefit from feature selection.

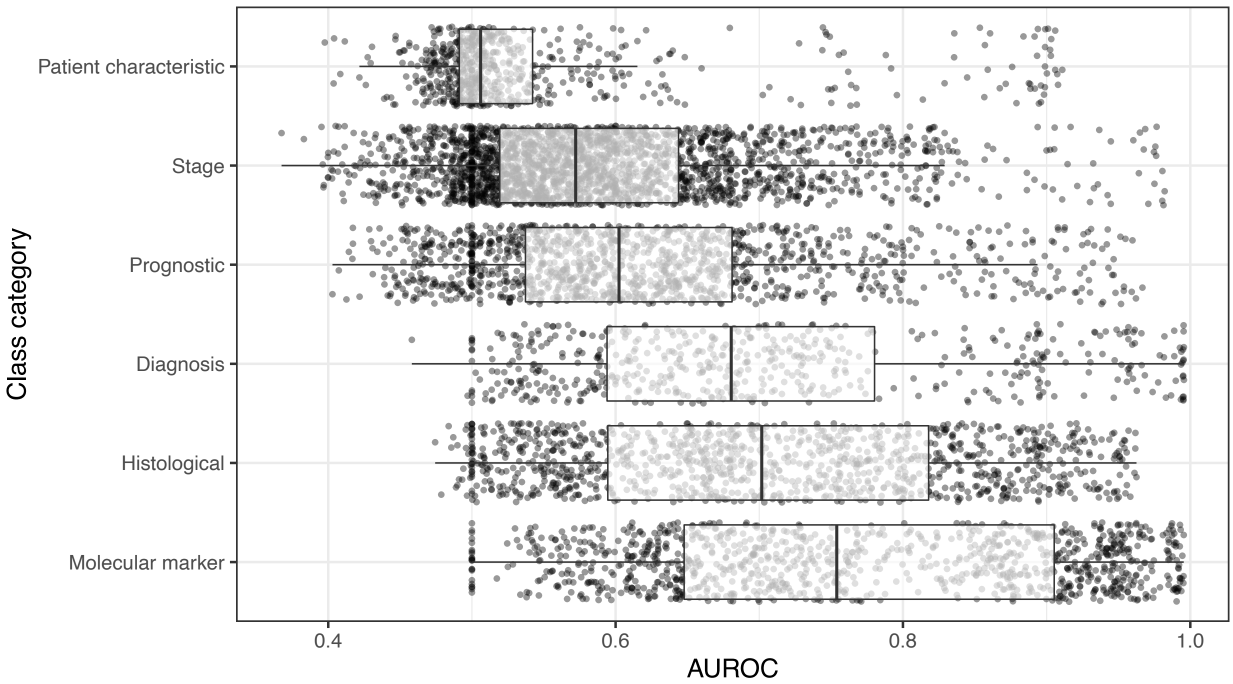


**Figure S21: Dataset performance by class category when using gene-expression and clinical predictors and performing feature selection.** For each class variable across all datasets, we assigned a category representing the type of patient state being predicted. For Analysis 5, we show the predictive performance for each combination of dataset, class variable, and classification algorithm in each class category. We use area under the receiver operating characteristic curve (AUROC) as a metric. The dashed, red line indicates the performance expected by random chance. The results are similar to those of Analyses 3 and 4 (Figures S9, S12).

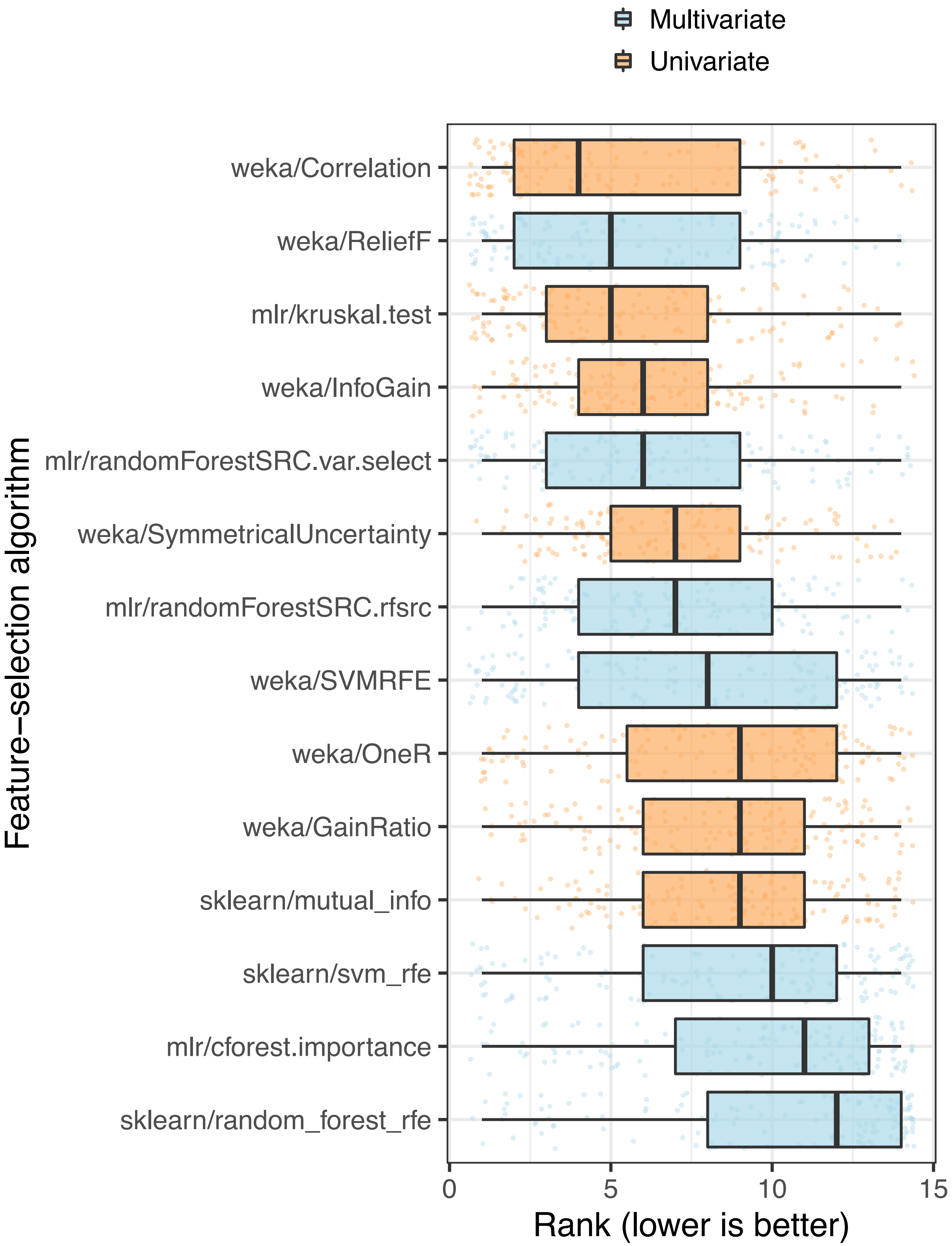
Multivariate
Univariate
Feature−selection algorithm
weka/Correlation
weka/ReliefF
mlr/kruskal.test
weka/InfoGain
mlr/randomForestSRC.var.select
weka/SymmetricalUncertainty
mlr/randomForestSRC.rfsrc
weka/SVMRFE
weka/OneR
weka/GainRatio
sklearn/mutual_info
sklearn/svm_rfe
mlr/cforest.importance
sklearn/random_forest_rfe
0
5
10
15
Rank (lower is better)

**Figure S22: Relative performance of feature-selection algorithms.** For Analysis 5, we used nested cross validation to estimate which features would be most informative for each algorithm in each training set. For each combination of dataset, class variable, and classification algorithm, we ranked the performance of the feature-selection algorithms based on area under the receiver operating characteristic curve (AUROC) and averaged the rankings across 5 iterations of Monte Carlo cross-validation. Each data point that overlays the box plots represents a particular dataset/class combination. Relatvely low average ranks are considered optimial. The `weka/Correlation` feature-selection algorithm performed best overall.

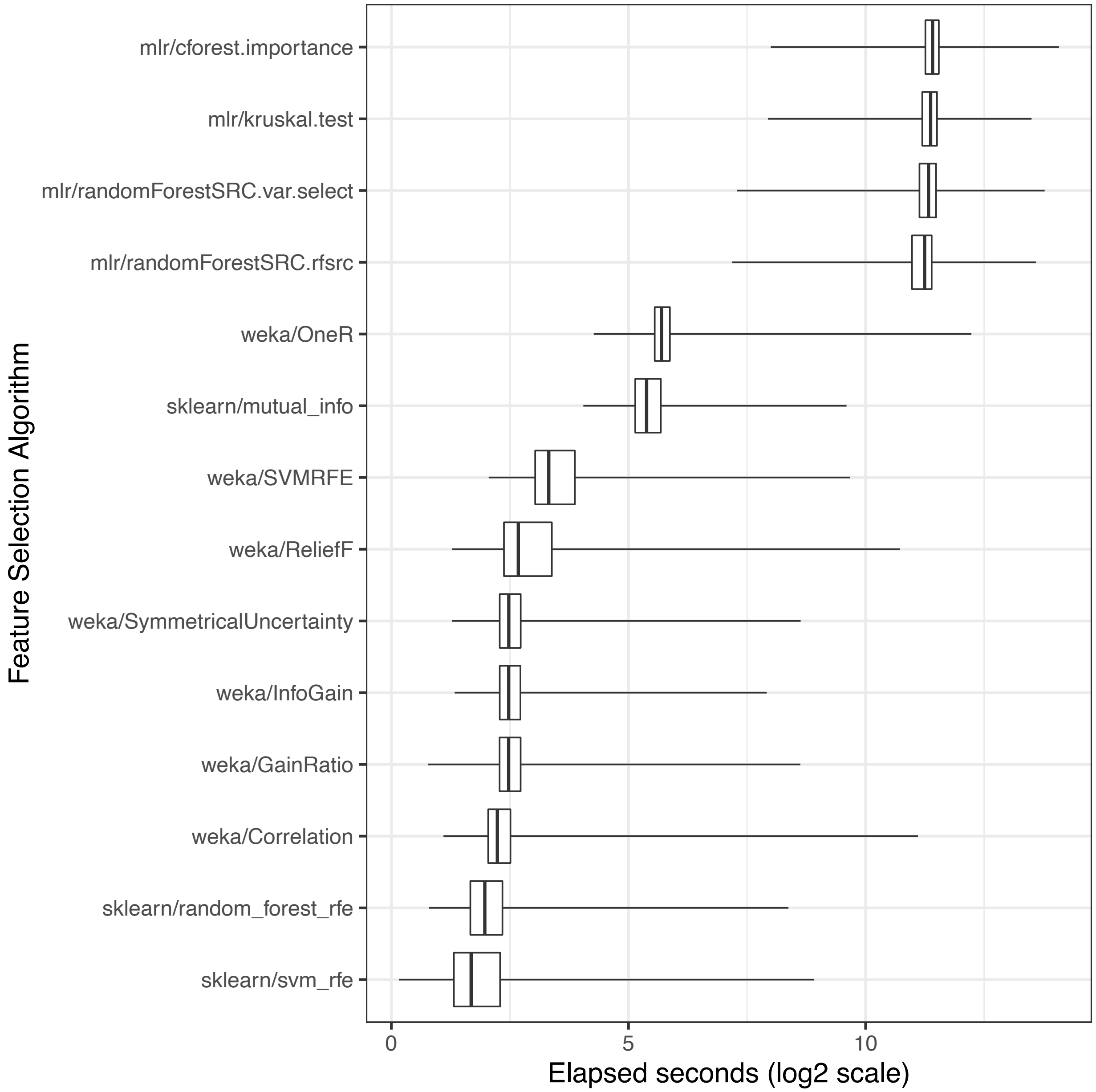


**Figure S23: Execution time per feature-selection algorithm.** In Analysis 5, we used nested cross validation to estimate which features were most informative for each training set. We calculated the time (in seconds) required by each feature-selection algorithm to rank the features. Then we averaged these times across all combinations of dataset, class variable, classification algorithm, and (outer) Monte Carlo cross-validation iteration. Some feature-selection algorithms were much more computationally intensive than others.

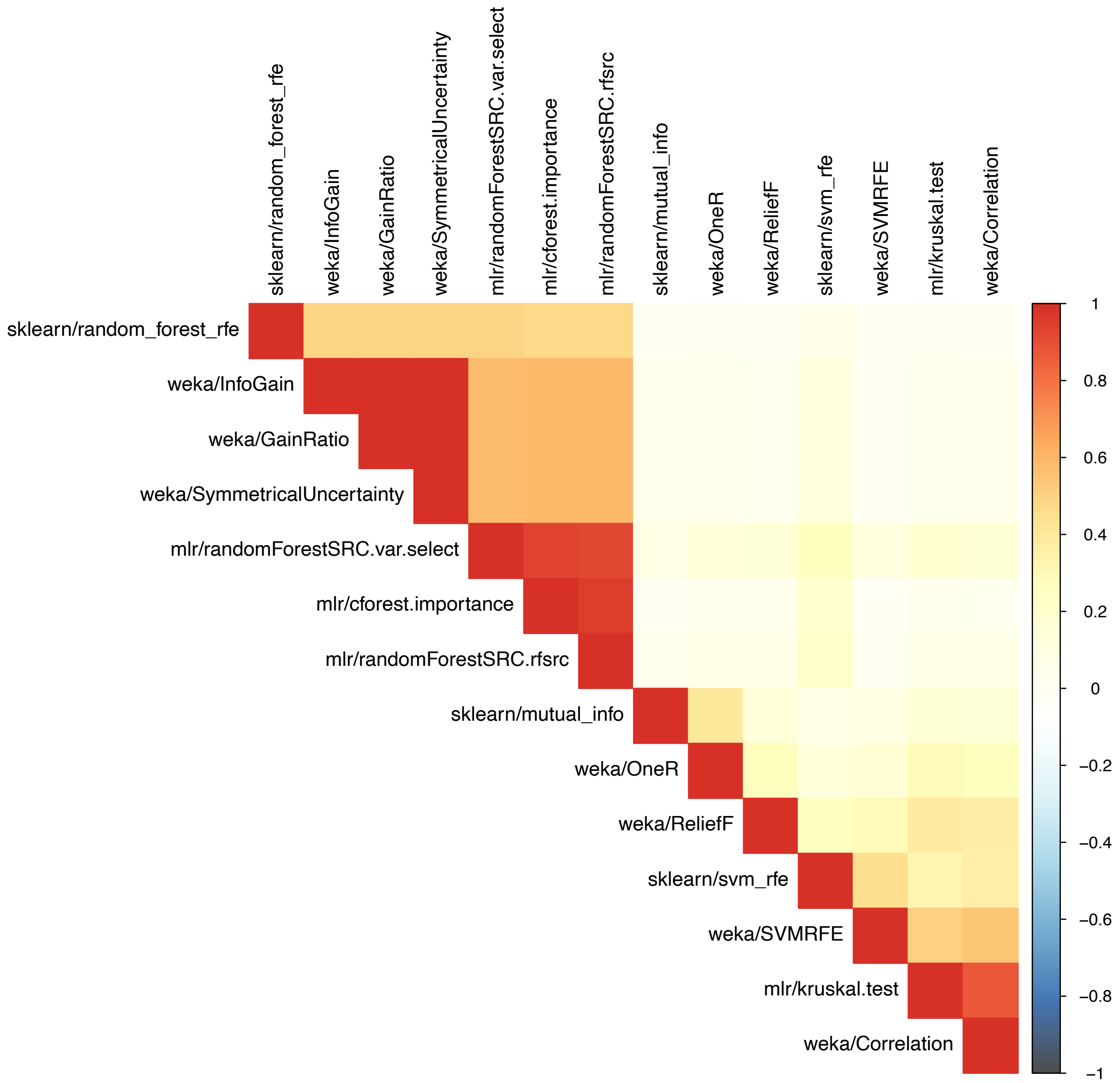


**Figure S24: Pairwise correlations of feature ranks between feature-selection algorithms for dataset GSE10320.** We used each feature-selection algorithm to rank the genes based on their informativeness for discriminating between relapse and non-relapse outcomes in Wilms tumor patients (GSE10320). After averaging the ranks across cross-validation iterations, we calculated the Spearman correlation coefficient for the feature ranks produced by each pair of algorithms. These coefficients are illustrated as a correlation plot.

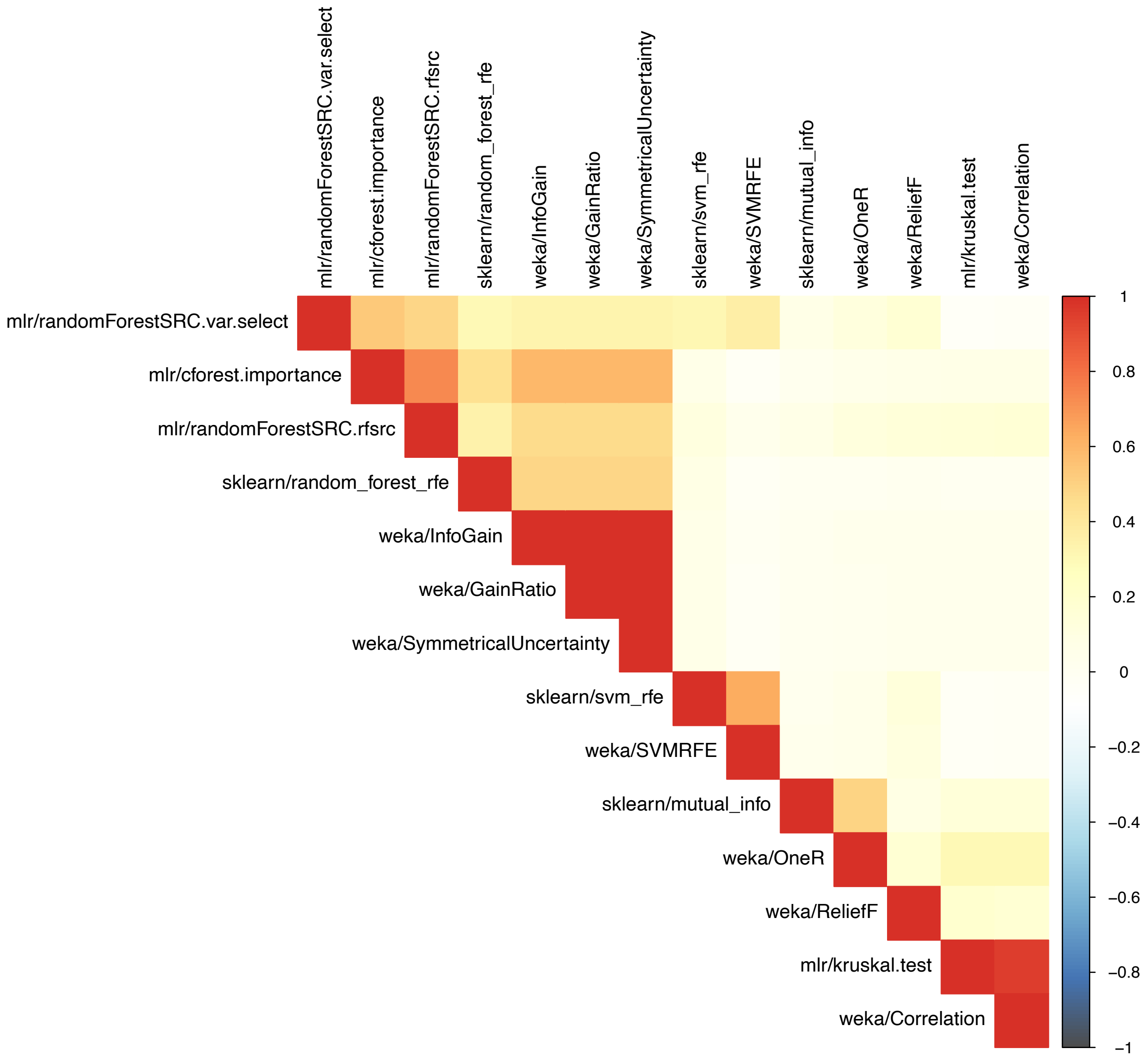


**Figure S25: Pairwise correlations of feature ranks between feature-selection algorithms for dataset GSE46691.** We used each feature-selection algorithm to rank the genes based on their informativeness for predicting early metastasis following radical prostatectomy (GSE46691). After averaging the ranks across cross-validation iterations, we calculated the Spearman correlation coefficient for the feature ranks produced by each pair of algorithms. These coefficients are illustrated as a correlation plot.

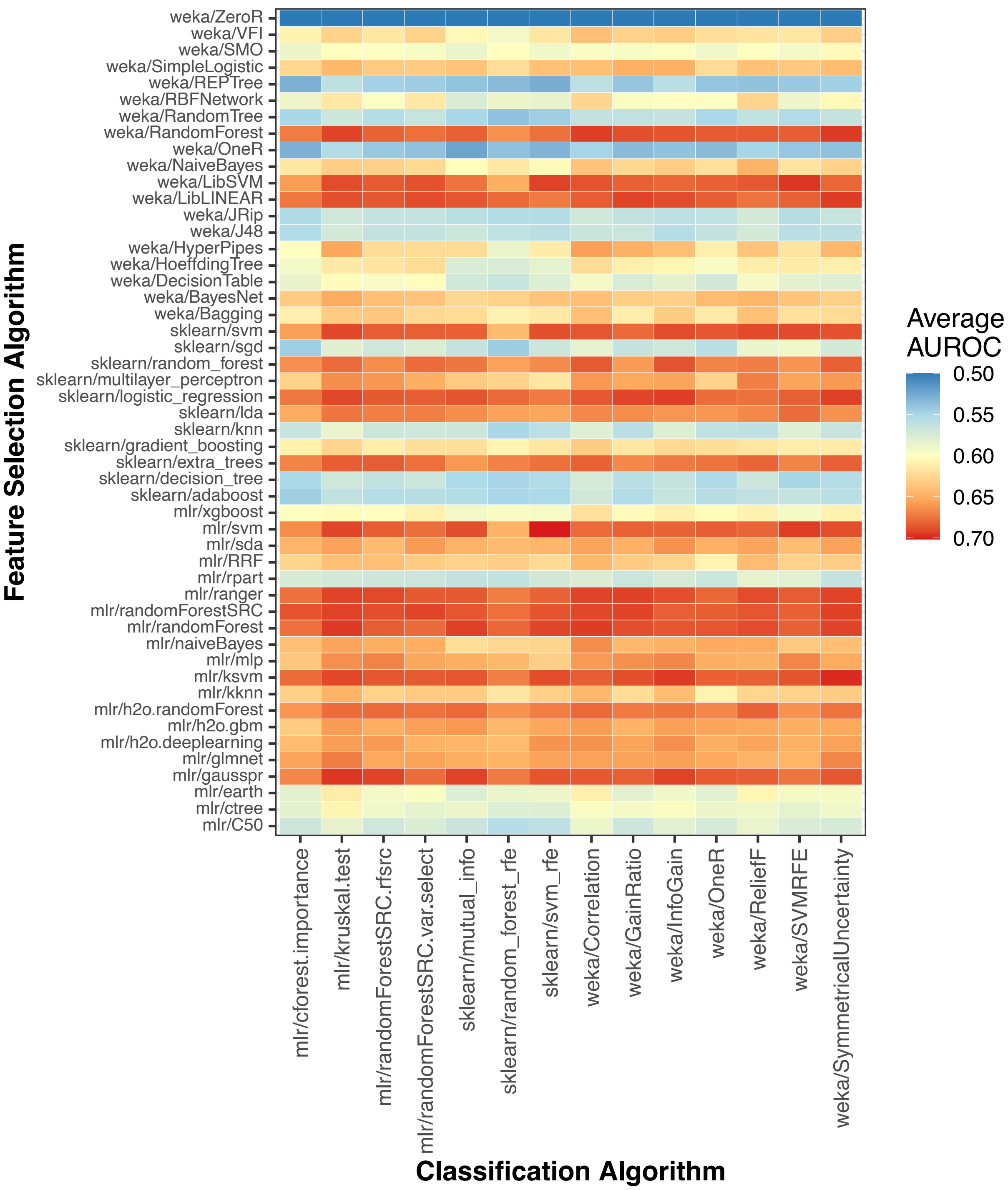


**Figure S26: Absolute classification performance per combination of feature-selection and classification algorithm.** For each combination of dataset and class variable, we averaged the area under the receiver operating characteristic curve (AUROC) across all Monte Carlo cross-validation iterations.

Then for each combination of feature-selection algorithm and classification algorithm, we calculated the median AUROC across all datasets and class variables.

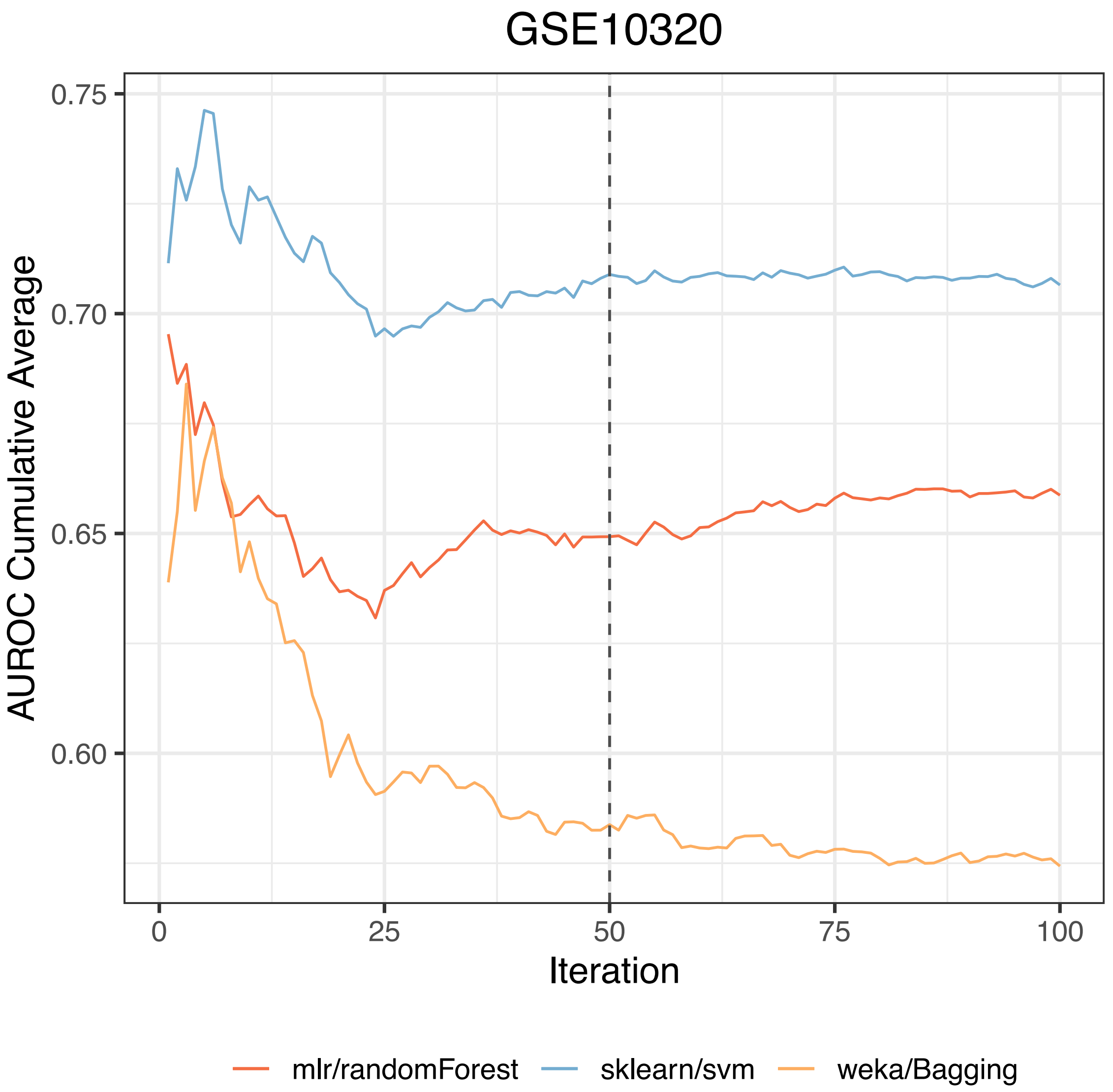


**Figure S27: Stability of classification performance for increasing numbers of cross-validation iterations on dataset GSE10320.** When using gene-expression predictors (Analysis 1), we estimated the number of Monte Carlo cross-validation iterations that would be sufficient to characterize algorithm performance. For three classification algorithms, we executed 100 cross-validation iterations on dataset GSE10320 (predicting relapse vs. non-relapse for Wilms tumor patients). As the number of iterations increased, we calculated the cumulative average of the area under the receiver operating characteristic curve (AUROC) for each algorithm. After performing at most 40 iterations, the cumulative averages did not change more than 0.01 over sequences of 10 iterations.

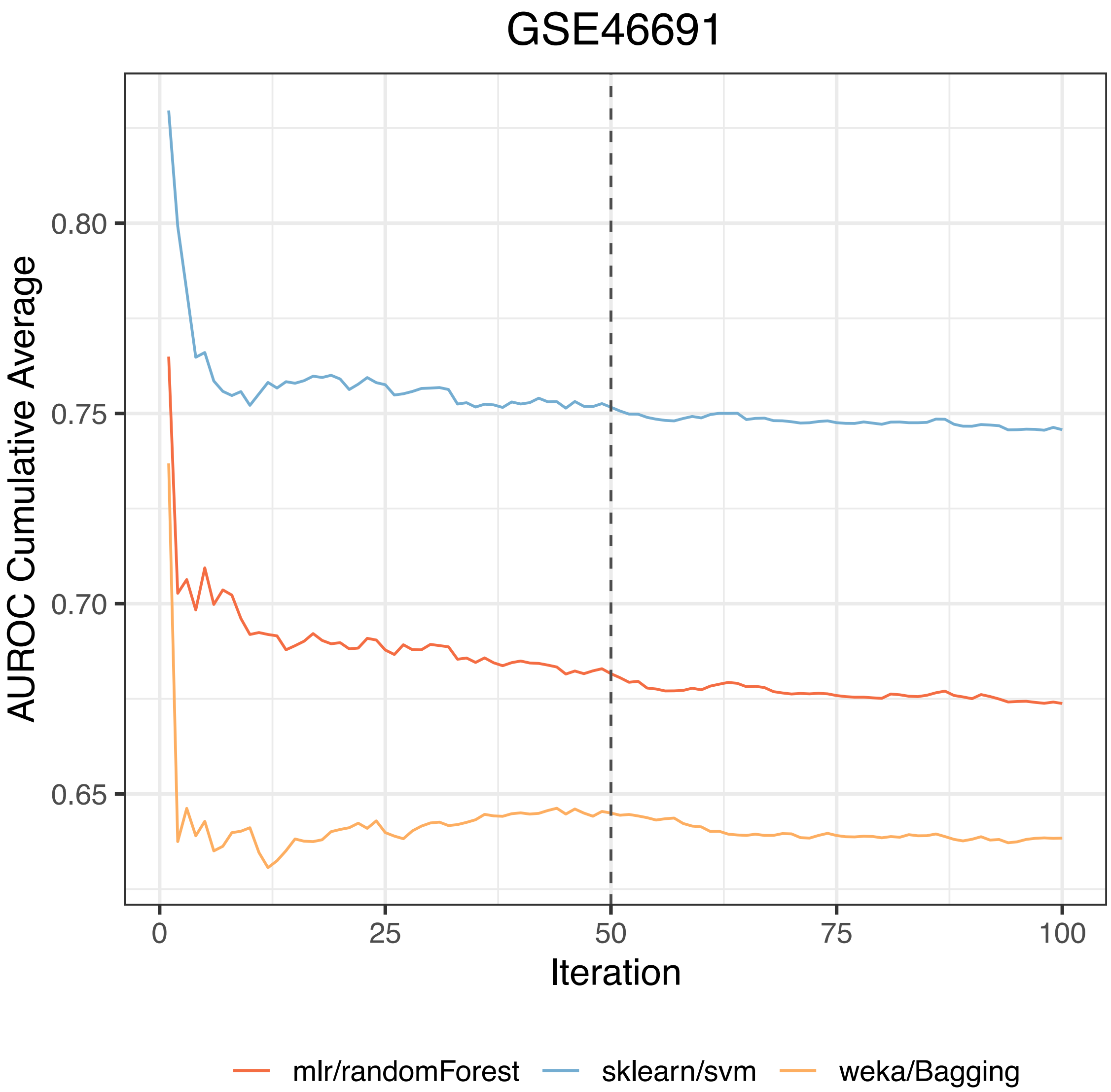


**Figure S28: Stability of classification performance for increasing numbers of cross-validation iterations on dataset GSE46691.** When using gene-expression predictors (Analysis 1), we estimated the number of Monte Carlo cross-validation iterations that would be sufficient to characterize algorithm performance. For three classification algorithms, we executed 100 cross-validation iterations on dataset GSE46691 (predicting early metastasis following radical prostatectomy). As the number of iterations increased, we calculated the cumulative average of the area under the receiver operating characteristic curve (AUROC) for each algorithm. After performing at most 22 iterations, the cumulative averages did not change more than 0.01 over sequences of 10 iterations.
